# Skeletal Muscle TFEB Signaling Promotes Central Nervous System Function and Reduces Neuroinflammation during Aging and Neurodegenerative Disease

**DOI:** 10.1101/2022.03.07.482891

**Authors:** Ian Matthews, Allison Birnbaum, Anastasia Gromova, Amy W. Huang, Kailin Liu, Eleanor A. Liu, Kristen Coutinho, Megan McGraw, Dalton C. Patterson, Macy T. Banks, Amber C. Nobles, Nhat Nguyen, Gennifer E. Merrihew, Lu Wang, Eric Baeuerle, Elizabeth Fernandez, Nicolas Musi, Michael J. MacCoss, Helen C. Miranda, Albert R. La Spada, Constanza J. Cortes

**Author notes:** **Funding**: NIH R01 AG077536 (CJC), University of Washington Nathan Shock Center P30 AG013280 (Pilot Grant to CJC), San Antonio Nathan Shock Center of Excellence in the Basic Biology of Aging P30AG013319 (Pilot Grant to CJC), NIH R35 122140 (ALS).

## Abstract

Skeletal muscle has recently arisen as a novel regulator of Central Nervous System (CNS) function and aging, secreting bioactive molecules known as myokines with metabolism-modifying functions in targeted tissues, including the CNS. Here we report the generation of a novel transgenic mouse with enhanced skeletal muscle lysosomal and mitochondrial function via targeted overexpression of Transcription Factor E-B (TFEB). We have discovered that the resulting geroprotective effects in skeletal muscle reduce neuroinflammation and the accumulation of tau-associated pathological hallmarks in a mouse model of tau pathology. Muscle-specific TFEB overexpression also significantly ameliorates proteotoxicity, reduces neuroinflammation and promotes transcriptional remodeling of the aged CNS, preserving cognition and memory in aged mice. Our results implicate the maintenance of skeletal muscle function throughout aging to direct regulation of CNS health and disease, and suggest that skeletal-muscle originating factors may act as novel therapeutic targets against age-associated neurodegenerative disorders.

## INTRODUCTION

Aging is associated with an organism-wide progressive loss of tissue form and function, broadly characterized by the ‘hallmarks of aging’ ^1^. In particular, the aging central nervous system (CNS) ^1,2^ exhibits a global loss in protein homeostasis (proteostasis), impaired neuroplasticity and resilience, and an increase in neuroinflammation ^2,3^. These alterations are believed to render the aging CNS vulnerable to age-associated dysfunction and the development of neurodegenerative disease ^2^.

Over the last ten years, there has been growing evidence that suggests prominent contributions of the periphery to the etiology of age-associated neurodegenerative diseases ^4–8^. In particular, manipulation of skeletal muscle protein quality control pathways yields important benefits to the invertebrate CNS, protecting against the accumulation of aggregation-prone neurodegenerative disease proteins in the brain and the retina ^5,8,9^. Although the mechanisms responsible for these benefits remain poorly understood, some of these effects are mediated by secreted factors that communicate metabolic and inflammatory signals between tissues ^5,7,9^. In agreement with this, blood circulating factors have recently arisen as potent regulators of mammalian CNS aging and metabolism ^10–13^. For example, exposure to a young plasma environment (either through heterochronic parabiosis or plasma transfers) can rescue function in the aging CNS ^12–14^ by decreasing neuroinflammation ^15^ and enhancing neurogenesis ^10^. Studies have further demonstrated that increasing the levels of these individual factors in peripheral circulation is sufficient to rejuvenate the aged CNS ^12,16,17^, supporting the existence of geroprotective circulating factors with CNS targeting effects. Although the source and identity of these neuroprotective circulating cytokines are unclear, several of them are expressed in, and are known to be secreted from, skeletal muscle ^5,18–20^. In fact, skeletal muscle acts as an endocrine organ, secreting a myriad of bioactive factors that induce metabolic changes in a non-cell autonomous manner in distant tissues like liver ^21^, adipose tissue ^22^, and the CNS ^19^.

Recently, this muscle-to-brain signaling axis has been proposed to have important implications for CNS aging and age-associated neurodegenerative disease ^19,23,24^. Indeed, skeletal muscle function has arisen as a key predictor for phenotypic and clinical outcomes in age-associated neurodegenerative diseases, including Alzheimer’s Disease (AD) and Parkinson’s Disease (PD) ^25,26^. Consistent with this, we were the first to uncover a novel mechanistic basis for motor neuron disease in the polyglutamine disease Spinal and Bulbar Muscular Atrophy (SBMA), where disruption of skeletal muscle autophagy initiates the pathogenic cascade that culminates in neuronal toxicity and death ^27^. Furthermore, we demonstrated that rescuing proteotoxicity specifically in skeletal muscle was sufficient to rescue motor neuron degeneration and extend lifespan in this model, despite robust misfolded protein burden in the brain ^28^. We also demonstrated the feasibility and efficacy of peripheral delivery of antisense oligonucleotides reducing skeletal muscle expression of the misfolded protein in rescuing neurological phenotypes ^29^, confirming that therapeutic targeting of skeletal muscle can have important benefits to the CNS.

Skeletal muscle metabolism is regulated in part by Transcription Factor E-B (TFEB), a master regulator of a lysosomal-to-nucleus signaling axis that integrates cellular metabolism and lysosomal function ^30–32^. TFEB expression and function are strongly induced in skeletal muscle in response to interventions with documented neuroprotective effects on the aging ^33,34^ and neurodegenerative disease-afflicted CNS, including low nutrient conditions ^35^ and exercise ^36^. Indeed, TFEB controls metabolic flexibility in muscle during exercise, inducing the expression of genes involved in mitochondrial biogenesis, fatty acid oxidation, and oxidative phosphorylation ^36^. This coordinated action optimizes mitochondrial substrate utilization, enhancing ATP production and exercise capacity ^36^. These findings identify TFEB as a critical mediator of the beneficial effects of exercise on metabolism. However, the precise role of TFEB-dependent signaling in skeletal muscle during aging, and its impact on the muscle-to-brain axis, remains largely unknown.

Here we report the generation of a novel transgenic mouse with enhanced muscle metabolism via overexpression of TFEB. We have discovered that the resulting enhanced skeletal muscle lysosomal function is maintained throughout the lifespan, protecting against the onset of age-associated mitochondrial dysfunction in aging skeletal muscle. Furthermore, we report that overexpression of TFEB in skeletal muscle significantly reduces hippocampal accumulation of neuropathological hallmarks and reduces neuroinflammation in a mouse model of tau pathology, despite no exogenous activation of the transgene in the CNS. Muscle TFEB overexpression also ameliorates proteotoxicity, reduces neuroinflammation, and promotes transcriptional remodeling of the aged CNS without neurodegenerative disease burden, preserving cognitive performance in healthy aging mice. Our results implicate, for the first time in mammals, maintenance of skeletal muscle function to direct regulation of CNS health, and suggest that skeletal-muscle originating factors may act as novel therapeutic targets against age-associated neurodegenerative diseases.

## RESULTS

### TFEB overexpression maintains fiber type composition, function, and metabolism-associated networks in aging skeletal muscle

We derived a novel line of conditional transgenic mice carrying the β-actin promoter in combination with the CMV enhancer (CAGGS) and with a floxed 3x-FLAG-eGFP STOP cassette placed just 5’ to a 3x-FLAG-human *TFEB* transgene (i.e. fxSTOP-TFEB transgenic mice) (**Figure 1A**). In the presence of Cre-recombinase, the STOP codon cassette will be excised out, allowing expression of the human TFEB transgene, also known as a Cre-loxP ON system. To drive overexpression of TFEB specifically in skeletal muscle, we crossed male fxSTOP-TFEB mice with female Human Skeletal Actin (HSA)-Cre mice to achieve widespread expression of Cre-recombinase in cells of the myogenic lineage ^37^.

**Figure 1:**
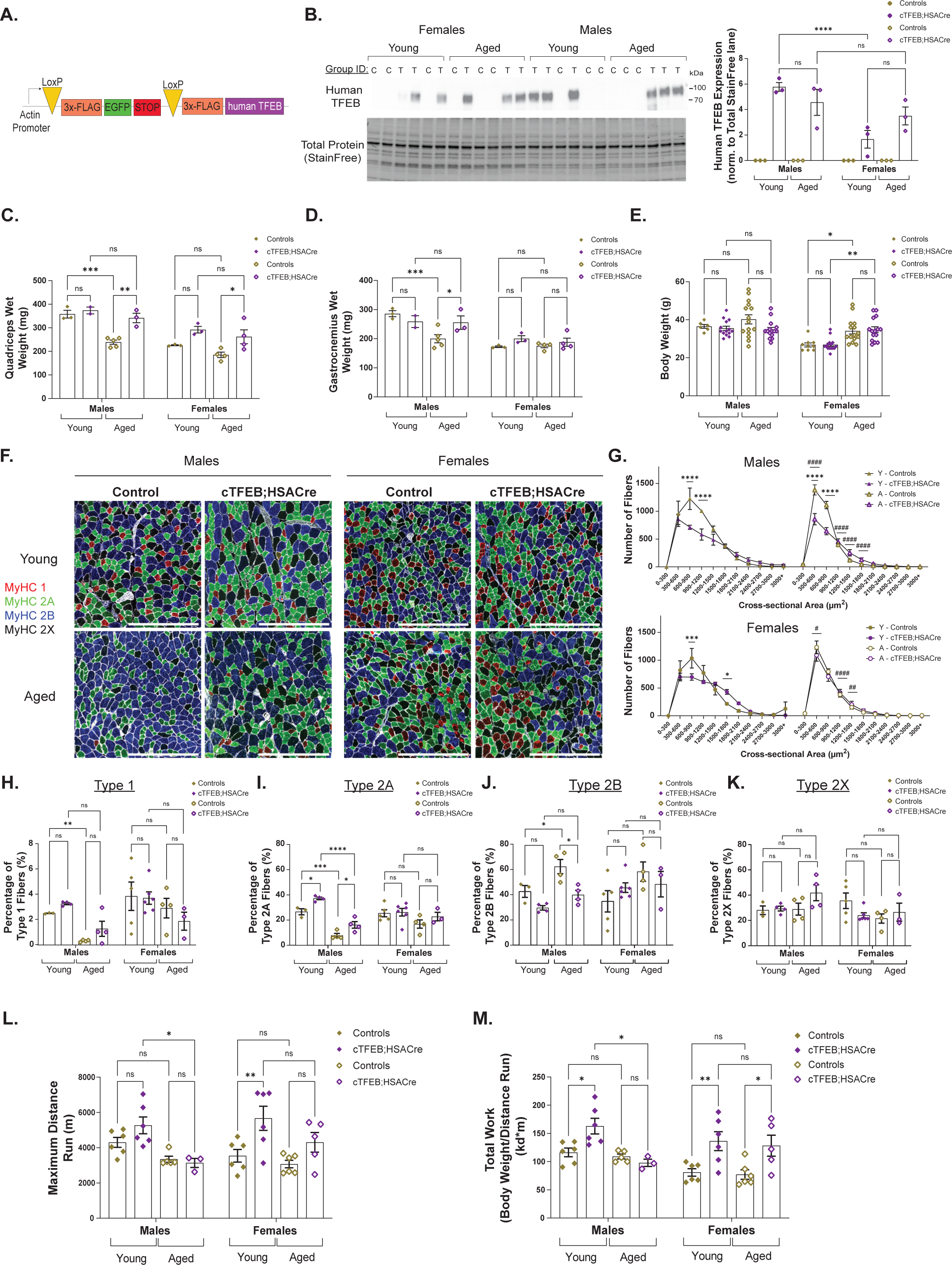
Skeletal muscle overexpression of TFEB preserves muscle size, fiber typing and increases exercise endurance during aging. (A) Schematic depicting 3x-FLAG human TFEB transgene. Cassettes are arranged from 5’ (left) to 3’ (right). Note that the 3x-FLAG-EGFP-STOP cassette (flanked by loxP sites) prevents expression of the 3x-FLAG human TFEB transgene unless in the presence of Cre recombinase. (B) Immunoblot of control and cTFEB;HSACre quadriceps lysates confirming 3x-FLAG-TFEB protein expression in young (5-6 months) and aged (22-24 months) mice of both sexes. C=control, T=cTFEB;HSACre individuals. Transgenic 3x-FLAG-TFEB protein weighs ∼75 kDa. Total Protein Stain Free lanes are shown below as a loading control. Densitometry quantification relative to average of young male control individuals to the right. (C-E) Skeletal muscle (quadriceps, C and gastrocnemius, D) wet weight and body weight (E) in young (6 months) and aged (22-24 months) control and cTFEB;HSACre transgenic mice of both sexes. (F) Representative images of fibertyping stains of cross-section of gastrocnemius muscle of young (7-9 months) and aged (22-24 months) control and cTFEB;HSACre transgenic mice of both sexes. MyHC1 (Fiber Type 1) in red, MyHC2A (Fiber Type 2A) in green, MyHC2B (Fiber Type 2B) in blue, MyHC2X (Fiber Type 2x) unstained, and laminin in white. Scale bars = 500 μm. (G) Fiber cross sectional area distribution curves from (F). *, statistical difference between control and cTFEB;HSACre. #, statistical difference between young and aged controls. Each data point represents the average of 3 separate images collected from 3 sections/individual. (H-K) Quantifications of each fiber type percentage from (F). Each data point represents the average of 3 separate images collected from 3 sections/individual. (L-M) High-intensity exhaustive treadmill exercise of young (6 months) and aged (18 months) control and cTFEB;HSACre transgenic mice of both sexes showing maximum distance run till exhaustion (L) and total work (body weight/distance run, M). Each point represents the average of all data from a single individual. Controls are age-matched littermates. Unless otherwise stated, data are presented as mean ± SEM. */^#^ p<0.05, **/^##^ p<0.01, ****/^####^ p<0.0001 ANOVA and post-hoc multiple comparisons for each sex. Lack of annotation indicates comparisons were not significant.

The resulting cTFEB;HSACre bigenic mice of both sexes exhibited a strong induction of human *TFEB mRNA* expression in skeletal muscle tissue (**Supplemental Figure 1A, quadriceps**). This exogenous *TFEB* expression was still prominent at 24 months of age in cTFEB;HSACre transgenic skeletal muscle (**Supplemental Figure 1B, quadriceps)** suggesting that overexpression of the transgene remains active even at very old ages. Immunoblotting of quadricep skeletal muscle confirmed robust expression of 3x-FLAG-TFEB protein in cTFEB;HSACre bigenic muscle lysates of both sexes (**Figure 1B**). Human TFEB protein levels in male cTFEB;HSACre mice amounted to about 2-fold higher levels than their female littermates at both ages examined. Importantly, and in agreement with our qRT-PCR studies, human TFEB was still robustly expressed at 24 months of age in both sexes, suggesting life-long expression of our transgene upon embryonic recombination (**Figure 1B**). As TFEB is a transcription factor, we confirmed nuclear localization of human TFEB in cTFEB;HSA-Cre skeletal muscle myonuclei via immunofluorescence (**Supplemental Figure 1C, in white**).

Next, we confirmed TFEB protein overexpression exclusively in skeletal muscle of cTFEB;HSACre bigenic mice by multiple approaches. First, we confirmed expression of the biologically inert 3x-FLAG-eGFP cassette upstream of the STOP codon (**Figure 1A**) and no exogenous expression of 3x-FLAG-TFEB in other highly metabolic tissues of non-myogenic lineages in cTFEB;HSACre transgenic mice, including brown adipose tissue (**Supplemental Figure 1D**) and liver (**Supplemental Figure 1E**). We detected a relatively mild expression of human *TFEB* mRNA levels in the heart of cTFEB;HSACre transgenic mice, confirming previous reports indicating low-level cross-over into cardiac tissue in the HSA-Cre model. However, this expression amounted to about ∼400-fold less in young animals and ∼15-fold less in aged animals than that detected in skeletal muscle at the same ages (**Supplemental Figure 1A, ventricles**). Importantly, we could not detect 3x-FLAG-TFEB protein via immunoblotting in cTFEB;HSACre heart ventricle lysates (**Supplemental Figure 1F**). This suggests that despite some HSA-Cre ‘leakiness’ into cardiac muscle, synthesis of exogenous TFEB protein in heart tissue is below detectable levels, confirming the high skeletal muscle tropism of HSA-Cre driven expression. Finally, we also did not detect expression of exogenous human TFEB mRNA in the hippocampus (**Supplemental Figure 1A & 1B, hippocampus**) or hemibrain protein lysates (**Supplemental Figure 1G**). To further confirm no CNS expression of transgenes associated with the HSA-Cre driver line, we directly assessed the Cre-mediated recombination of an additional transgene reporter in TdTomato Ai9;HSACre mice. We confirmed robust tdTomato fluorescence in skeletal muscle (**Supplemental Figure 1H**) but no detectable TdTomato fluorescence in the CNS (**Supplemental Figure 1I**), including the hippocampus (**Supplemental Figure 1I, insets**), of TdTomato Ai9;HSACre mice, consistent with the muscle-restricted specificity of this HSA-Cre-loxP-ON approach ^38^. Overall, these data confirm previous reports of striated muscle-specific Cre-mediated recombination using the HSA-Cre transgenic driver line ^37^, and demonstrate detectable expression of human TFEB protein only in skeletal muscle of cTFEB;HSACre bigenic mice. This establishes our newly developed inducible fxSTOP-TFEB mouse as an effective model to generate spatiotemporally controlled transgenic expression of human TFEB.

Analysis of skeletal muscle wet weight, a proxy for skeletal muscle size, revealed trends towards larger muscle wet weights in young cTFEB;HSACre animals, and a robust and significant preservation of quadriceps and gastrocnemius skeletal muscle wet weight in aged (22–24 month-old) transgenic mice of both sexes (**Figure 1C-D**). Indeed, while control animals lost around 20-30% of their quadriceps wet weight during aging, there was no significant differences between young or aged cTFEB;HSACre skeletal muscle weight (**Figure 1C-D**). Interestingly, this maintenance of muscle wet weight was not associated with changes in total body weight at either age examined (**Figure 1E**). Consistent with these findings, we determined that overexpression of TFEB significantly decreases the number of small fibers (<1500 µm^2^) and increases the number of large fibers (>1500 µm^2^) in young gastrocnemius skeletal muscle of both sexes (**Figure 1F-G**). This was accompanied by an increase in abundance of type 1 (∼1% more) and type 2A (∼9% more) fibers (**Figure 1H-K**) and an increase in the cross-sectional area (CSA) of Type 2B (∼200 um^2^ larger) and 2X (∼300 um^2^ larger) fibers with no changes in Feret’s minimum diameter (**Supplemental Figure 2A-H)**, which was prevalent in male mice and trending in female mice. Importantly, while control male gastrocnemius muscle lost a significant portion of their large fibers during aging, they were not altered in aged male cTFEB:HSACre transgenic muscle (**Figure 1F-G**). The abundance of Type 1 and Type 2A fibers was also still prevalent in aged male cTFEB:HSACre transgenic muscle (**Figure 1H-K and Supplemental Figure 2G-H)**, suggesting that TFEB overexpression remodels both fast-twitch glycolytic and oxidative fiber fibers and that these changes are maintained during aging. To determine the physiological effects of this fiber type switch on skeletal muscle function, we examined exercise performance in all of our cohorts. We determined that at 6 months of age, muscle-TFEB overexpressing mice had a 20% (males) and 40% (females) longer maximum running distance to exhaustion (**Figure 1L**) and engaged in more total work (**Figure 1M**) than their control littermates. Importantly, this increased enhanced physical performance was still present in aged female cTFEB;HSACre transgenic mice (**Figure 1M**). Taken together, these results suggest that lifelong-expression of TFEB prevents fiber type atrophy and alters fiber type switching in aging skeletal muscle, ultimately preserving muscle size and running performance during aging.

Given the principal role of TFEB-regulated signaling (i.e. lysosomal and mitochondrial function) to skeletal muscle health ^39,40^, as well as the profound geroprotective effects of TFEB overexpression on muscle fiber types, muscle size and exercise performance in aging muscle (**Figure 1**), we sought to examine the overall changes in the skeletal muscle protein landscape in response to TFEB overexpression throughout aging. We performed proteome profiling of young and aged, control and cTFEB;HSACre male quadricep skeletal muscle. Differential abundance analysis of species revealed 909 proteins associated with TFEB overexpression in young (6-month-old) skeletal muscle (**Figure 2A**). Of these, 857 (∼94%) were significantly overrepresented and 52 (∼6%) were underrepresented compared to their littermate controls. At 18 months of age, a time point where age-associated dysfunction is prevalent in murine skeletal muscle ^41^, the same proteomics approach identified 351 differentially expressed proteins in TFEB-expressing skeletal muscle (**Figure 2B**). Of these, 283 proteins (∼81%) were significantly enriched and 68 proteins (∼19%) were significantly decreased in cTFEB;HSACre bigenic muscle, with only 53 of these proteins being uniquely expressed in older mice. Importantly, at both ages examined, TFEB remained significantly overexpressed (**Figure** 2A & 2B**, boxes**), confirming our previous finding that exogenous skeletal muscle TFEB expression is maintained across the lifespan of our cTFEB;HSACre model (**Figure 1B**).

**Figure 2:**
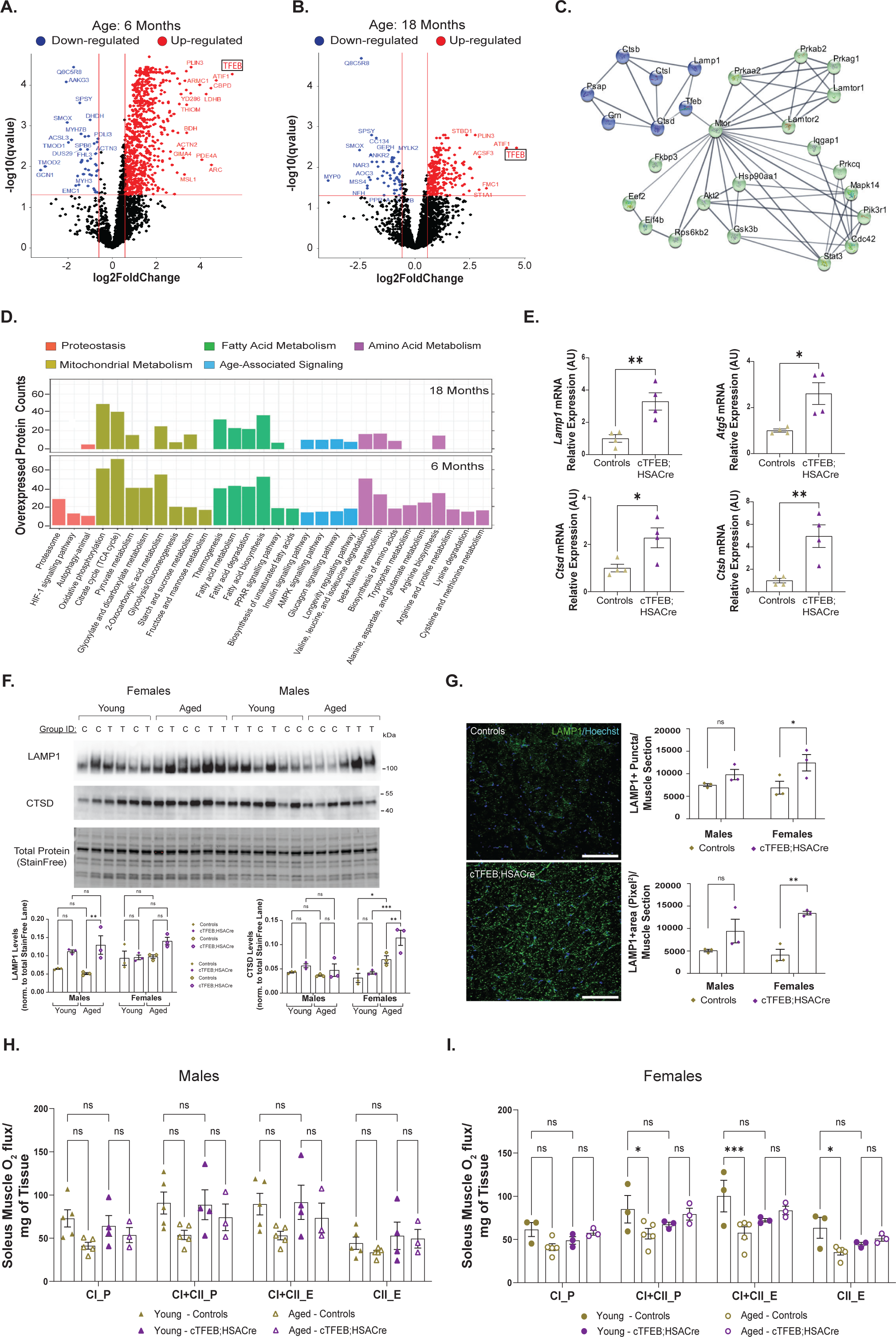
Muscle TFEB increases lysosomal network size and preserves mitochondrial function during aging. (A-B) Volcano plot of proteomics analysis depicting differentially regulated proteins in young (A, 6 months old) and aged (B, 18 months old) cTFEB;HSACre quadriceps muscle relative to controls. Overexpressed proteins in red, underexpressed proteins in blue. TFEB (rectangles) is highly expressed in both aged and young muscle. (C) STRING predicted protein-protein interactions of overexpressed proteins from (A) showing interacting networks converging on TFEB (blue) and mTOR (green). (D) KEGG enrichment analysis clusters overexpressed proteins from (A-B) into categories associated with proteostasis and autophagy (orange), mitochondrial metabolism (gold), fatty acid metabolism (green), age-associated signaling (blue), and amino acid metabolism (purple). (E) qRT-PCR for *Lamp1*, *Atg5*, *Ctsd*, and *CtsB* expression in quadriceps mRNA lysates from young (6 months old) male cTFEB;HSACre mice and controls. (F) Immunoblot analysis for LAMP1 and Cathepsin D protein from young (5-6 months) and aged (22-24 months) control and cTFEB;HSACre quadriceps lysates mice of both sexes. Total Protein Stain Free lanes are shown as a loading control. Marker densitometry quantification relative to Protein Stain Free densitometry shown below. (G) Representative images of gastrocnemius cross section from young (6-7 months old) control and cTFEB;HSACre female mice stained for LAMP1 (green), nuclei/Hoechst (blue). Quantification of LAMP1+ puncta and LAMP1+ area/muscle section for both sexes are shown on the right. Scale bars = 100 µm. (H-I) Mitochondrial function measured by oxygen flux normalized to citrate synthase activity in soleus muscle from young (6 months old) and aged (18 months old) male (H) and female (I) control and cTFEB;HSACre semi-permeabilized soleus muscle. CI_P is complex I and CI+CII_P is complex I and II linked respiration coupled to ATP production. CI+CII_E is uncoupled complex I and II respiration or maximum respiration. CII_E is complex II respiration in the uncoupled state. Each point represents the average of all data collected from an individual. Controls are age-matched littermates. All data were analyzed by 2-way between-subject (age × genotype) ANOVA with post hoc multiple comparisons, or one-way ANOVA and multiple comparisons post-hoc for each sex. Data are presented as mean ± SEM. * p<0.05, ** p<0.01. Lack of annotation indicates comparisons were not significant.

To visualize the key networks of differentially expressed proteins associated with TFEB-overexpression, we performed a functional enrichment analysis on all overexpressed proteins using the STRING protein-protein interaction network ^42^ (**Figure 2C**). Using TFEB as our ‘bait’ for functional connectivity nodes, we identified two clear interaction networks: the first one was associated with lysosomal biogenesis and function (**Figure 2C, blue**), and included Lamp1 (an essential lysosomal structural protein), Grn (progranulin) and Psap (prosaposin) as well as the lysosomal proteases: Ctsb (Cathepsin B), Ctsd (Cathepsin D), and Ctsl (Cathepsin L). The second interaction network is associated with key skeletal muscle proteostasis and metabolic signaling pathways including mTOR, Akt and GSK3β (**Figure 2C, green**). Both of these nodes are key elements of the signaling network controlled by TFEB-mediated transcription ^31,32^, were previously reported to be transcriptionally elevated in skeletal muscle after viral-mediated TFEB overexpression ^36^, and are central regulators of aging ^1^.

To further understand the functions of all the upregulated proteins in TFEB-overexpressing skeletal muscle, KEGG enrichment analysis was performed. As expected, central proteostasis categories were significantly upregulated in young TFEB-overexpressing skeletal muscle: including Hypoxia-inducible factor 1 (HIF-1) signaling, proteasome, and autophagy/lysosomal function categories (**Figure 2D, orange**). Importantly, this enrichment in proteostasis categories and TFEB-regulated nodes of signaling was also present in our aging cohorts (**Figure 2D, top**). We confirmed functional activation of TFEB-dependent transcription in cTFEB:HSA-Cre bigenic muscle, using our previously reported TFEB-responsive muscle gene targets ^27^ including *Lamp1*, *Atg5* (a key regulator of autophagy initiation), *Ctsd* and *Ctsb* (**Figure 2E**).

Immunoblotting studies confirmed significant increases in Lamp1 and Ctsd protein in cTFEB;HSACre bigenic muscle (**Figure 2F**). Consistent with previous studies demonstrating TFEB overexpression increases the number and size of LAMP1-positive lysosomes in multiple tissues ^30,35,43^, we also detected significant increases in the number of LAMP1+ puncta and total LAMP1+ area in cross-sections of young female (and similar trends in male) cTFEB;HSACre skeletal muscle (**Figure 2G**). There were no significant changes to LC3II/LCI ratios or total polyubiquitinated protein levels in cTFEB;HSACre transgenic mice (**Supplemental Figure 2I**). These findings confirm physiologically significant increases in TFEB-dependent transcription ^36^ and lysosomal biogenesis ^32,36^ in skeletal muscle of cTFEB:HSA-Cre mice.

Additional functional categories enriched in our proteomics analysis related to cellular metabolism and mitochondrial function, including thermogenesis, oxidative phosphorylation and pyruvate metabolism (**Figure 2D, gold**). We also detected central fatty acid metabolism pathways, including PPAR signaling and metabolism, and degradation and biosynthesis of fatty acids (**Figure 2D, green**). This is consistent with previously published reports suggesting that acute TFEB expression in young skeletal muscle induces the expression of genes involved in mitochondrial biogenesis and oxidative phosphorylation ^36^. Here we confirm for the first time that these changes in cTFEB;HSACre muscle are maintained throughout aging, particularly those associated with mitochondrial metabolism and fatty acid metabolism (**Figure 2D, top, gold and green**).

We assessed whether these proteomic profiles associated with mitochondrial function translated into mitochondrial function differences using high-resolution respirometry with simultaneous measurements ROS production assays ^44^. As reported previously for soleus skeletal muscle ^44^, oxygen consumption rates decrease with aging across all mitochondrial complexes in semi-permeabilized soleus muscle of control mice of both sexes (6 vs. 18 months of age, gold icons) (**Figure 2H-2I**). Whereas there was no difference in mitochondrial respiration rates between control and TFEB-overexpressing skeletal muscle at young ages (closed icons), respiration rates of aged cTFEB;HSACre transgenic muscle (open icons) remained comparable to their young counterparts across all mitochondrial complexes examined (**Figure2 H-2I**), particularly in female muscle. Together with our functional data showing increases in exercise capacity in cTFEB;HSACre mice (**Figure 1L-M**), these results suggest that muscle-TFEB overexpression promotes maintenance of mitochondrial function and bioenergetic reserves, preventing the age-associated decline of mitochondrial respiration commonly observed in aging skeletal muscle ^1,44^.

Finally, proteomics KEGG analysis also showed enrichment in cTFEB;HSACre bigenic muscle for classical signaling pathways shown to decline during aging, including the Insulin-IGF1-signaling (IIS) and AMPK signaling ^1^ (**Figure 2D, blue**). We also detected multiple hits for amino acid metabolism pathways, such as catabolism of branched-chain amino acids (BCAAs), including valine, leucine and isoleucine metabolism (**Figure 2D, purple**). Importantly, these changes were still evident in our aged cohorts (**Figure 2D, top**), consistent with our functional studies suggesting that TFEB overexpression throughout the lifespan has multiple benefits in skeletal muscle, targeting established hallmarks of aging ^1^.

The skeletal muscle ‘secretome’ (the totality of released organic and inorganic molecules released from muscle resident cells) is highly dynamic, responding to physiological and pathophysiological stimuli ^45,46^ and potentially also changing with age ^46^. To directly examine any changes to the skeletal muscle protein-based secretome associated with TFEB-overexpression, we pursued additional *in silico* analysis through the Vertebrate Secretome Database (VerSeDa) on our proteomics cohorts to identify unique age- and genotype-associated signatures of potentially secreted proteins ^47^ differentially expressed in cTFEB;HSACre skeletal muscle. We identified multiple proteins with predicted secreted profiles to be enriched in young and aged TFEB-expressing skeletal muscle (**Figure 3A**). Some secreted proteins exhibited upregulation (e.g., nucleobindin-1, angiopoietin-related protein 1) or down-regulation (collagen alpha-1 (I) chain), only in young cTFEB;HSACre muscle (**Figure 3A, green**). Others remained up-regulated at both ages in cTFEB;HSACre muscle, (prosaposin, mammalian ependymin-related protein 1 (MERP1) and dehydrogenase/ reductase SDR family member 7C, D/R SDR7C) (**Figure 3A, overlap**), or were only differentially expressed in aged muscle (biglycan-1, PG-S1, **Figure 3A, blue**), highlighting the dynamic nature of the skeletal muscle secretome throughout aging. Cathepsin B (Ctsb), a previously documented muscle-originating secreted factor with known CNS-targeting effects ^20^ and a known target of TFEB-dependent transcription (**Figure 2E**) ^32^ was our top up-regulated hit at both ages examined. We validated significant increases in pro-enzyme levels of Ctsb in both sexes and at both ages examined (**Figure 3B**), as well as significant increases in mature (secreted) Ctsb isoforms in aged cTFEB;HSACre transgenic muscle (**Figure 3B**). We also detected trends towards increased levels of circulating CTSB in the serum of female cTFEB;HSACre mice (**Figure 3E**). Given the potent remodeling of the lysosomal network associated with activation of TFEB-mediated transcription (**Figure 2**) ^30–32^, our data suggests that overexpression of TFEB in skeletal muscle drives increased expression and secretion of mature Ctsb, a CNS targeting-factor, into circulation.

**Figure 3:**
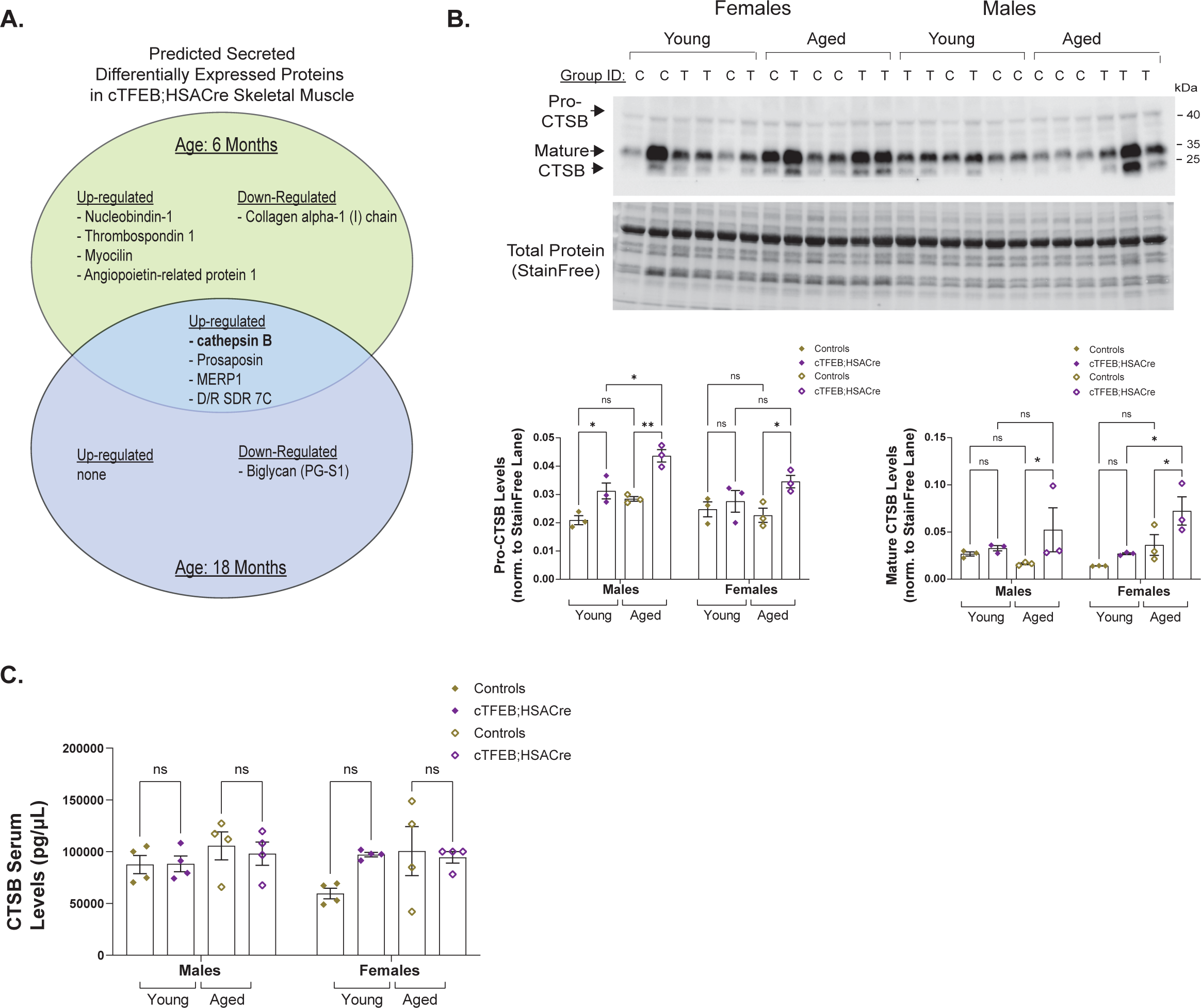
Increased levels of known CNS-targeting myokines in TFEB-expressing skeletal muscle. (A) VerSeDa-predicted secreted proteins identified as up-regulated or down-regulated in young (green) and aged (blue) cTFEB;HSACre transgenic muscle, from proteomics studies from Figure 2A-B. (B) Immunoblot analysis for Cathepsin B protein in quadriceps protein lysates from young (5-6 months) and aged (22-24 months) control and cTFEB;HSACre quadriceps lysates mice of both sexes. Note that the mature fragment of Cathepsin B is the secreted form. Total Protein Stain Free lanes are shown as a loading control. Marker densitometry quantification relative to Protein Stain Free densitometry shown below. (C) Levels of cathepsin B in serum from young (5-6 months) and aged (22-24 months) control and cTFEB;HSACre mice of both sexes. Each point represents the average of technical replicates from an individual. Controls are age-matched littermates. All data were analyzed by 2-way between-subject (age × genotype) ANOVA with post hoc multiple comparisons, data are presented as mean ± SEM. * p<0.05, ** p<0.01. Lack of annotation indicates comparisons were not significant.

### Enhanced skeletal muscle TFEB-signaling reduces accumulation of hyperphosphorylated tau and microglial activation in a mouse model of tau pathology

To assess any potential neuroprotective effects of enhancing skeletal muscle TFEB-signaling in the context of age-associated neurodegenerative disease pathologies, we derived cTFEB;HSACre transgenic mice on the MAPT P301S (PS19) background. This is a well-known model of neurofibrillary tangle toxicity, a hallmark of AD and related tauopathies ^48^, and is characterized by prominent accumulation of hippocampal hyperphosphorylated human mutant MAPT protein and pronounced neuroinflammation, including microgliosis and astrocyte reactivity ^48^.

We first confirmed muscle-restricted 3x-FLAG-TFEB overexpression in skeletal muscle lysates from MAPT P301S;cTFEB;HSACre mice, as well as the presence of human Tau in hippocampal lysates of both groups via immunoblotting (**Figure 4A**). As previously reported ^48^, at 9 months of age there is robust accumulation of Ser202/Thr205 hyperphosphorylated tau (residues abnormally phosphorylated in paired helical filaments during AD progression) in the dentate gyrus of the hippocampus of MAPT P301S mice of both sexes ^48^ (**Figure 4B, middle panel**). Conversely, we noted a significant reduction in the total fluorescence counts of AT8-positive phosphorylated tau in the dentate gyrus of MAPT P301S;cTFEB;HSACre mice (**Figure 4B**, **purple**) compared with MAPT P301S littermate controls (**Figure 4B**, **teal**). More so, this reduction was particularly evident in the levels of ‘intracellular tau’ accumulating around hippocampal cell bodies (**Figure 4B, insets**), suggesting reduced accumulation of toxic tau species and aggregates ^49^ in the hippocampus of MAPT P301S;cTFEB;HSACre mice. We confirmed similar but less striking reductions in Thr231 phospho-tau (another primary site of tau phosphorylation associated with neurotoxic tau accumulation) immunostaining (**Supplemental Figure 3A**), suggesting overall reductions in tau hyperphosphorylation in the hippocampus of MAPT P301S transgenic mice with TFEB overexpression in skeletal muscle.

**Figure 4:**
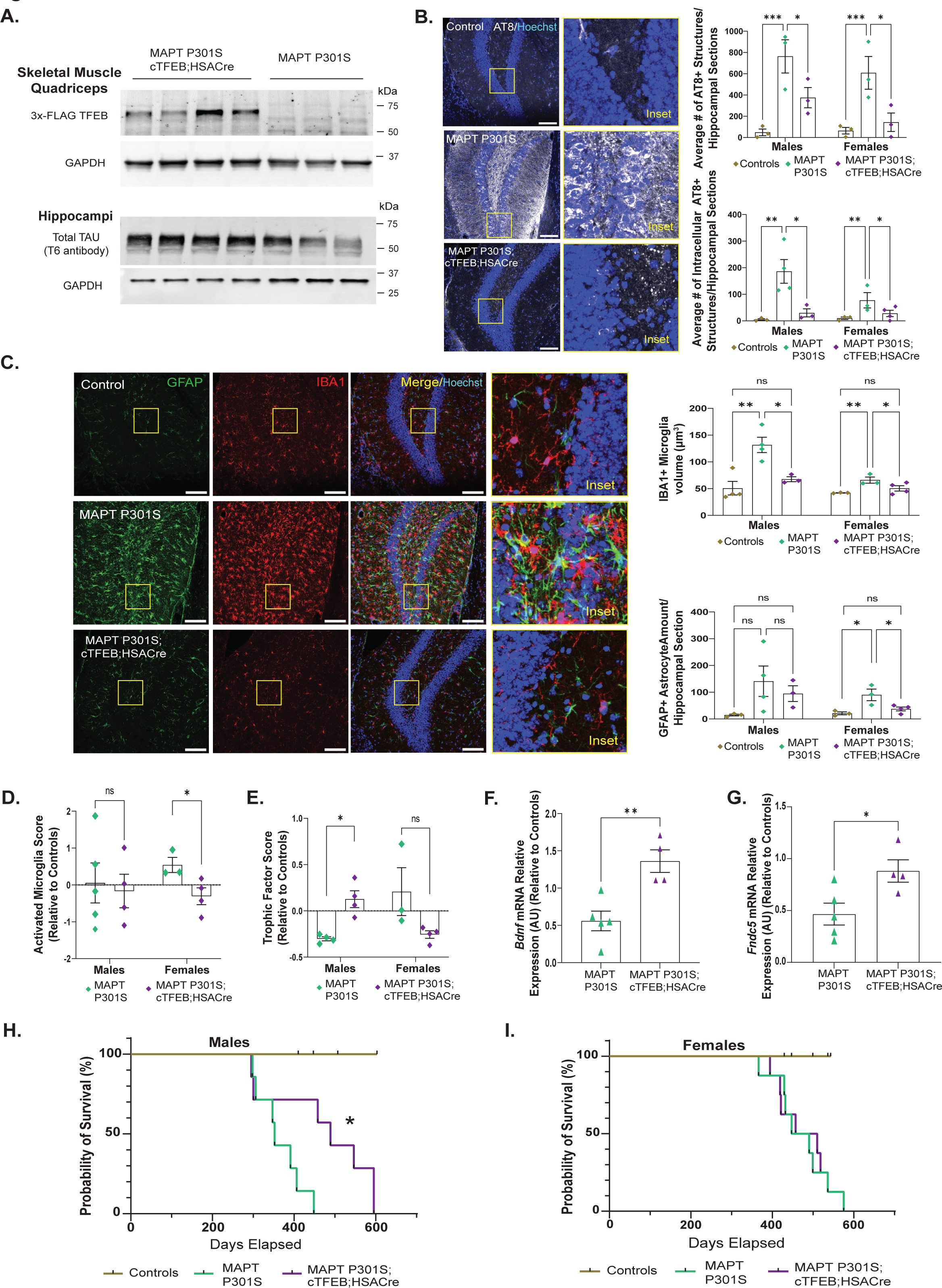
Skeletal muscle TFEB-overexpression reduces accumulation of hyperphosphorylated tau and microglial activation in a mouse model of tau pathology. (A) Immunoblot of 3x-FLAG TFEB protein in quadriceps muscle lysates from 9-month old MAPT P301S;cTFEB;HSACre and MAPT P301S mice (top). Immunoblot for total Tau protein (T6 antibody) in hippocampal lysates from same individuals (bottom). GAPDH is shown below as a loading control. (B) Representative merged images of the hippocampus dentate gyrus stained for phosphorylated tau (AT8, white) and nuclei/Hoechst (blue) in 9-month old control (top, in gold), MAPT P301S (middle, in teal), and MAPT P301S;cTFEB;HSACre mice (bottom, in purple). Insets depicting perinuclear AT8 staining are 5X zooms of areas demarcated by yellow squares. Representative images are from male mice. Quantification of total phosphotau staining/section (right, top) and for intracellular phosphotau staining, (right, bottom). Each point represents the average of all sections containing the dentate gyrus (2-3 sections) for an individual animal. Scale bars = 100 µm. (C) Representative merged images of the dentate gyrus stained for astrocytes (GFAP, green), microglia (IBA1, red) and nuclei (Hoechst, blue) in 9-month old control (top, gold), MAPT P301S (middle, teal), and MAPT P301S;cTFEB;HSACre mice (bottom, purple). Insets depicting glia morphology are 5X zooms of areas demarcated by yellow squares. Representative images are from male mice. Quantification of IBA1+ microglia volume (top, right) and GFAP+ astrocyte staining amounts per section (bottom, right). Each point represents the average of all sections containing the dentate gyrus (2-3 sections) for an individual animal. Scale bars = 100 µm. (D-E) Nanostring nCounter AD panel cluster analysis of differentially expressed genes for gene clusters associated with microglial activation (D) and with trophic factors (E) in P301S MAPT hippocampi (teal) compared to MAPT P301S;cTFEB;HSACre (purple) age-matched littermates of both sexes. Analysis done via nSolver (Nanostring) differential gene expression analysis software. Each point represents cluster scoring for a single individual hippocampal mRNA lysate. (F-G) qRT-PCR for *Bdnf* (F) and *Fndc5* (G) expression in hippocampal RNA lysates from 9-month old male MAPT P301S (teal) and MAPT P301S;cTFEB;HSACre mice (purple) relative to controls (not shown). (H-I) Survival curves for male (H) and female (I) control (gold), MAPT P301S (teal), and MAPT P301S;cTFEB;HSACre mice (purple). Controls are littermates. Unless otherwise noted, data are represented as mean ± SEM. * p<0.05, ** p<0.01,*** p<0.001, One-way ANOVA, post-hoc multiple comparisons (B,C), t-test (D-G) and log-rank (Mantel-Cox) test (H-I). Lack of annotation indicates comparisons were not significant.

Astrocytes and microglia are key mediators of neuroinflammation, and undergo key morphological and functional alterations during neurodegenerative disease, including changes in ramification and cellular process complexity ^50^, reduced immune surveillance activity ^51^ and increased reactivity phenotypes ^52^. We confirmed significant increases in the staining for GFAP-positive structures (a marker of astrocyte reactivity) in the dentate gyrus of 9-month-old MAPT P301S mice of both sexes (**Figure 4C, teal**) compared to their MAPT-negative littermate controls (**Figure 4C, gold**). We also detected significant increases in the volume of IBA1+-positive microglia in the same brain region (**Figure 4C, teal**), consistent with previous reports in the MAPT P301S model indicating high levels of neuroinflammation starting as early as 4 months of age ^48,53^. Strikingly, we noted significant reductions in both of these pro-inflammatory glial morphometric parameters in the hippocampus of MAPT P301S;cTFEB;HSACre littermates (**Figure 4C, purple**), suggesting overall reduced hippocampal neuroinflammation during symptomatic disease stages through activation of skeletal muscle TFEB-expression.

### Enhancing TFEB-signaling in skeletal muscle promotes neurotrophic signaling and reduces hippocampal neuroinflammation in MAPT P301S transgenic mice

To gain more precise insights into disease-relevant transcriptional changes in the hippocampus of MAPT P301S mice with enhanced muscle-TFEB expression, we used the Nanostring nCounter® Mouse Alzheimer’s Disease panel. This platform directly assesses the expression of 770 targeted genes curated from human and pre-clinical models of AD and robustly captures disease-relevant signatures and their modifications after pre-clinical interventions ^54^. Using this approach, we found 24 individual differentially expressed genes in MAPT P301S;cTFEB;HSACre hippocampi compared to their littermate MAPT P301S+ controls between both sexes (**Supplemental Figure 3B-C**). Interestingly, most differentially regulated genes were downregulated (18 of 24), consistent with their identity as transcriptional drivers of disease ^54^. Many of these transcripts are associated with microglial activation (*Rhoc, Clic1)* markers of neuronal function (*Gdap1l1, Amot, Map3k9),* phospholipid and cholesterol remodeling (*Ano6, Cyp27a1*), growth factor signaling (*Vgf*), endothelial cell migration and matrix adhesion (*Rras, Cdc42Ep1*) and guanine nucleotide exchange factors (*Dock3*). Multiple down-regulated genes were associated with novel variants and loci associated with increased polygenic neurodegenerative disease risk (*Prkd3, Arhgap31)* as well as predictors of the rate of cognitive decline in AD (*Hecw1* ^55^). *Pbxip1,* another significantly downregulated marker, has been found to be associated with phosphorylated-tau and Aβ_1–40_ levels in the human temporal cortex^56^. Additional transcriptional changes of significance include genes of unknown function but are classified as transcriptional identifiers of AD progression *(Ptprn, Cpne2,* and *Eri2)*^54^. Up-regulated transcripts included gene repressors (*Zbtb33*), modulators of neuroinflammation (*Pecam1*) and neurodegeneration (*Scna*), as well as a carbohydrate transmembrane transporter (*Tmem144*) and a GTP-ase nucleotide exchange factor (*Rcc1*).

Annotation of the differentially co-expressed/co-functional clusters included in this panel revealed significant reductions in the nCounter AD microglial activation module (composed of 63 genes associated with microglial activation during AD progression) (**Figure 4D**) in the hippocampi of 6 month old female (but not male) MAPT P301S;cTFEB;HSACre mice compared with MAPT P301S littermates. This is consistent with reduced neuroinflammation in the hippocampus of MAPT P301S mice with enhanced muscle-TFEB expression, as suggested by our glial morphometric analysis (**Figure 4C**). Interestingly, we also detected a sex-bias in the trophic factor functional cluster (composed of genes *Arhgdib, Bcl2, Calm3, Camk4, Gab1, Mapk9, Mapkapk2, Ntrk2, Pik3r1* and *Psen1*), with a significant improvement back to baseline expression in the hippocampus of male (but not female) MAPT P301S;cTFEB;HSACre mice (**Figure 4E**). We confirmed activation of neurotrophic signaling as early as 6 months of age in the hippocampus of male MAPT P301S;cTFEB;HSACre mice by targeted qRT-PCR analysis, which showed corrections of declining expression levels of neurotrophic factors *Bdnf* and *Fndc5* back to control littermate levels (**Figure 4F-G**) ^23,57^. Importantly, there was also a significant extension (38%) in the lifespan of male MAPT P301S;cTFEB;HSACre mice (**Figure 4H**). While the median survival of male MAPT P301S mice was 352 days (**Figure 4H, teal curve**), male MAPT P301S;cTFEB;HSACre mice median survival was 489 days (**Figure 4H, purple curve).** Although we did not detect a similar lifespan extension in female MAPT P301S;cTFEB;HSACre mice (**Figure 4I**), it is worth noting that in our hands, female MAPT P301S transgenic mice already live significantly longer than their male littermate controls (median survival of 468.5 days), perhaps suggesting they have reached maximal lifespan in this model. Finally, we did not detect any differences in ataxia progression ^58^ (**Supplemental Figure 3D**) or weight loss (**Supplemental Figure 3E**) between our groups. Altogether, these provocative results suggest that enhanced skeletal muscle-TFEB expression modifies the accumulation of pathogenic tau isoforms, reduces neuroinflammation and re-activates neurotrophic signaling in the hippocampus of MAPT P301S mice in a sex-dependent manner. Importantly, this occurs in the absence of detectable exogenous human TFEB expression in the CNS (**Supplemental Figure 1**), suggesting an independent mechanism of muscle-to-brain communication underlying these effects.

### Skeletal muscle over-expression of TFEB decreases neuroinflammation markers and lipofuscin accumulation in the healthy aged CNS

To examine the effect of increased muscle-TFEB expression on the healthy aging brain without neurodegenerative disease burden, we assessed the transcriptional and functional status of known inflammatory markers in the CNS of healthy aged (20+ months old) control and cTFEB;HSACre mice, an age when global decreases in proteostasis and increases in pro-inflammatory signaling are detectable in most murine tissues ^1,59^, including the CNS ^1,2^. Total hippocampal mRNA levels of *Ccl2* and *NfKb*, pro-inflammatory cytokines previously reported to mediate microglial responses to inflammation, were significantly reduced in the hippocampus of aging female (but not male) cTFEB;HSACre bigenic mice (**Figure 5A-B**). Interestingly, the expression of *Il6* (interleukin 6) was significantly elevated in the same groups (**Figure 5C**). We also detected significantly higher levels of expression of *Bdnf* (Brain Derived Neurotrophic Factor) in hippocampal lysates from aged male (but not female) mice (**Figure 5D**), similar to the upregulation of hippocampal BDNF we observed in our male MAPT P301;cTFEB;HSACre model (**Figure 4F**).

**Figure 5:**
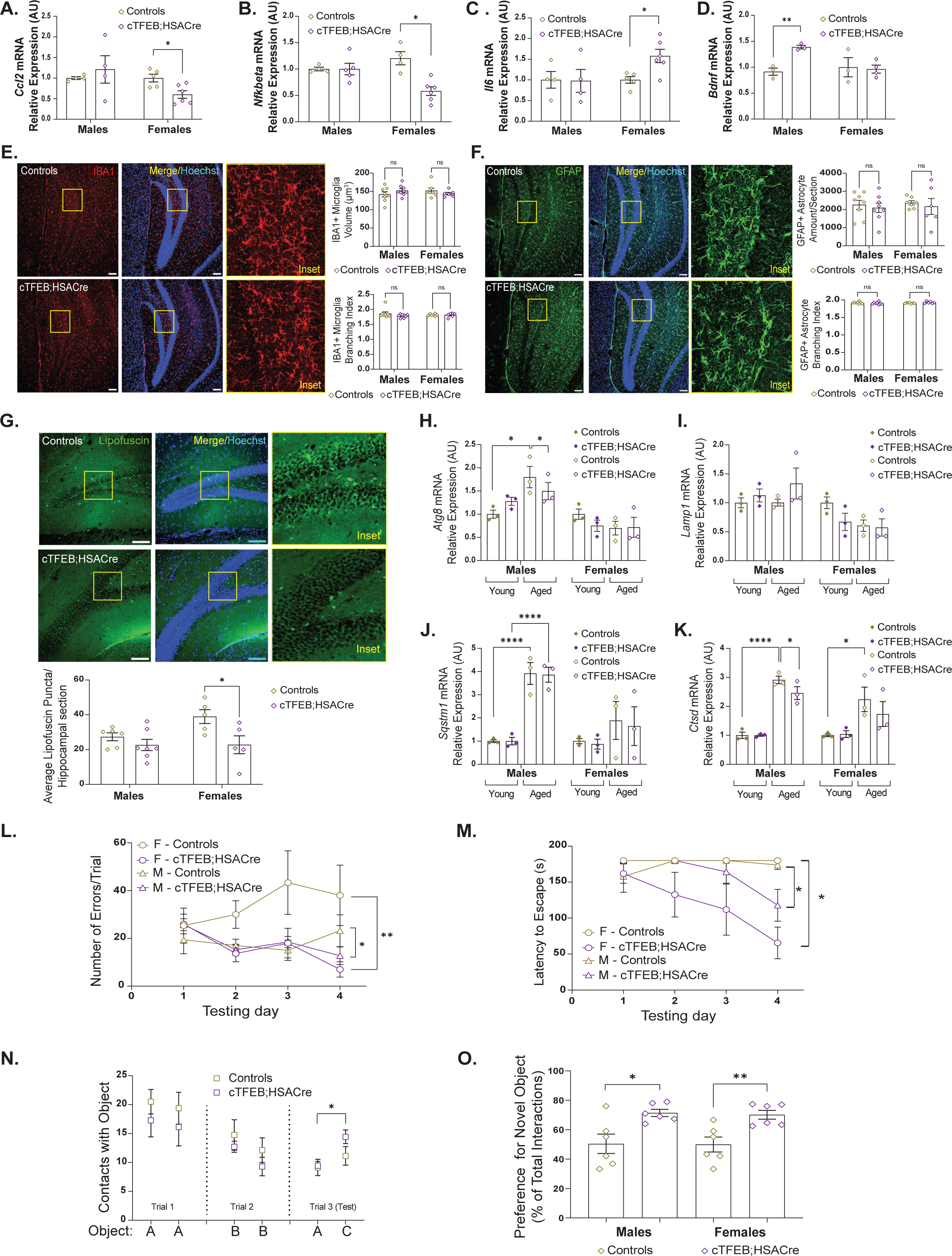
Skeletal muscle TFEB-overexpression decreases neuroinflammation, lipofuscin accumulation, and improves cognitive performance in aged mice. (A-D) qRT-PCR for *Ccl2* (A), *Nfkb* (B), *Il6* (C), and *Bdnf* (D) expression in hippocampal mRNA lysates from 18-month old cTFEB;HSACre mice and controls. (E-F) Representative images dentate gyrus of the hippocampus stained for Hoechst (blue), IBA1 (E, red, representative images are from females), and GFAP (F, green, representative images are from males) in 18-22-month old control (top) and cTFEB;HSACre (bottom) mice. Insets depicting glia morphology are 5X zooms of areas demarcated by yellow squares. Quantification of IBA1+ microglial volume (E, top right) and IBA1+ branching index (E, bottom right). Average number of GFAP+ astrocytes pixels per section (F, top) and GFAP+ astrocyte branching index (F, bottom). Each point represents the average of all sections containing the dentate gyrus from an individual. Scale bars = 100 µm. (G) Representative merged images of the dentate gyrus of 18-22-month old control (top) and cTFEB;HSACre (middle) mice stained for Hoechst (blue), and lipofuscin puncta (autofluorescence, green, representative images are from females). Insets depicting autofluorescent puncta within the dentate gyrus are 5X zooms of areas demarcated by yellow squares. Scale bars = 100 µm. (H-K) qRT-PCR for *Atg8* (H), *Lamp1* (I), *Sqstm1* (J), and *Ctsd* (K) expression in hippocampal lysates from young (6 months old, closed diamonds) and aged (21 months old, open diamonds) cTFEB;HSACre mice (purple diamonds) and controls (gold diamonds). (L-Q) Neurocognitive phenotyping of aged (18 months old) cTFEB;HSACre and control animals. Barnes maze (L,M), novel object recognition (N,O). Controls are age-matched littermates. Each point represents the average of all data collected from an individual. Data are represented as mean ± SEM. * p<0.05, ** p<0.01, One-way ANOVA or two-way RMANOVA, post-hoc-tests. Lack of annotation indicates comparisons were not significant.

Using similar morphometric analysis as before to quantify glial cell shape and morphology, we did not detect any differences in microglia (IBA1+) or astrocyte (GFAP+) number, volume or ramification state in the dentate gyrus of aged cTFEB;HSACre bigenic mice compared to their littermate controls (**Figure 5E-F**). This may suggest that the expression differences observed via transcriptional approaches above may either originate in non-glial cells (neurons or endothelial cells, for example) or not translate into detectable morphological changes at the ages examined. It is important to note that age-associated changes in neuroinflammation in an otherwise healthy brain are not as pronounced as those seen in the context of neurodegenerative disease (such as in the MAPT P301S model, for example). Overall, these results suggest a shift in the cytokine transcriptional landscape of the aging female hippocampus with upregulated skeletal muscle TFEB-expression towards reduced inflammatory signaling, consistent with what we observed in our MAPT P301S studies (**Figure 4**).

Accumulation of lipofuscin, a non-degradable intracellular auto-fluorescent polymer, becomes prominent in the aging brain, likely reflecting an age-associated decline in basal CNS autophagy ^1^. We examined the clearance capabilities of the aging CNS by indirect immunofluorescence of lipofuscin granules in brain sections of aged (21-24 months old) control and cTFEB;HSACre mice. While we detected striking lipofuscin granule accumulation in several brain regions of aged control mice, including the hippocampus (**Figure 5G, top**), there was a significant decrease in lipofuscin deposition in the dentate gyrus of aged cTFEB;HSACre females (**Figure 5G, bottom),** with a similar non-significant trend in males. To determine any changes in proteostasis associated signaling in this brain region, we next examined the transcriptional levels of multiple autophagy and lysosomal associated genes.

Consistent with previous reports, we confirmed significant age-associated increases in the expression of autophagy lysosomal markers *Atg8/Lc3*, *Lamp1, Sqstm1,* and *Ctsd* in the male hippocampus, which has been proposed to occur as a compensation for age-associated decline in proteostasis function (**Figure 5H-K**). Interestingly, these differences were not as prominent in the aging control female hippocampus. Furthermore, some of these transcriptional increases were reduced in the hippocampus of aged male cTFEB;HSACre bigenic mice, suggesting that enhancing skeletal muscle TFEB-signaling promotes maintenance of hippocampal proteostasis during aging. Our data thus indicate a potential sex-bias in activation of neuroprotective signaling in the hippocampus of aging cTFEB:HSACre bigenic mice, with females showing decreased neuroinflammation (**Figure 5A-C**) and males showing increased neurotrophin expression levels (**Figure 5D**). Altogether, our results suggest that overexpression of TFEB in skeletal muscle throughout the lifespan promotes maintenance of protein quality control regulation in the CNS, ultimately reducing proteostatic burden and neuroinflammation in the aging hippocampus, similar to what we observed in our MAPT P301S studies.

### Improved performance in neurocognitive testing of aging cTFEB;HSACre mice

We next pursued neurocognitive testing in aged (16-18 months old) control and cTFEB;HSACre mice. First, we evaluated spatial learning and memory using the Barnes maze task, a hippocampal working memory test known to be sensitive to aging in mice. We documented a significant decrease in the number of errors per trial (**Figure 5L**) and significantly faster escape times (**Figure 5M**) in aged cTFEB;HSACre mice of both sexes in comparison to controls. By the last trial, cTFEB;HSACre mice escaped the maze twice as quickly as control mice. In the novel object recognition task – an independent behavioral test of hippocampal recognition memory – we also found that aged cTFEB;HSACre mice of both sexes exhibited a significantly greater number of contacts with the novel object compared to controls during the test phase (**Figure 5N**). Indeed, aged cTFEB;HSACre mice have around a ∼15% higher preference for the novel object relative to their age-matched littermate controls (**Figure 5O**). Importantly, we also confirmed no differences in visual performance and motor activity (not shown) between aged control and cTFEB;HSACre cohorts. Hence, these results provide exciting evidence for a pronounced improvement in neural function in the aging brain of cTFEB;HSACre mice.

## Enhancing skeletal muscle TFEB-signaling transcriptionally remodels the hippocampus of cTFEB;HSACre bigenic mice throughout the lifespan in a sex-dependent manner

To determine the molecular basis of the benefits of enhancing skeletal muscle TFEB-signaling on the aging CNS, we performed unbiased transcriptome analysis on hippocampal RNAs isolated from young (6 months old) and aged (22-24 months old) mice (**Figures 6-7**). By this time point, aging control animals have increased neuroinflammation (**Figure 5A-B**), have evident CNS proteostasis dysfunction (**Figure 5G**), and display deficits in cognitive function (**Figure 5L-O**), phenotypes that are significantly improved in cTFEB;HSACre mice.

**Figure 6:**
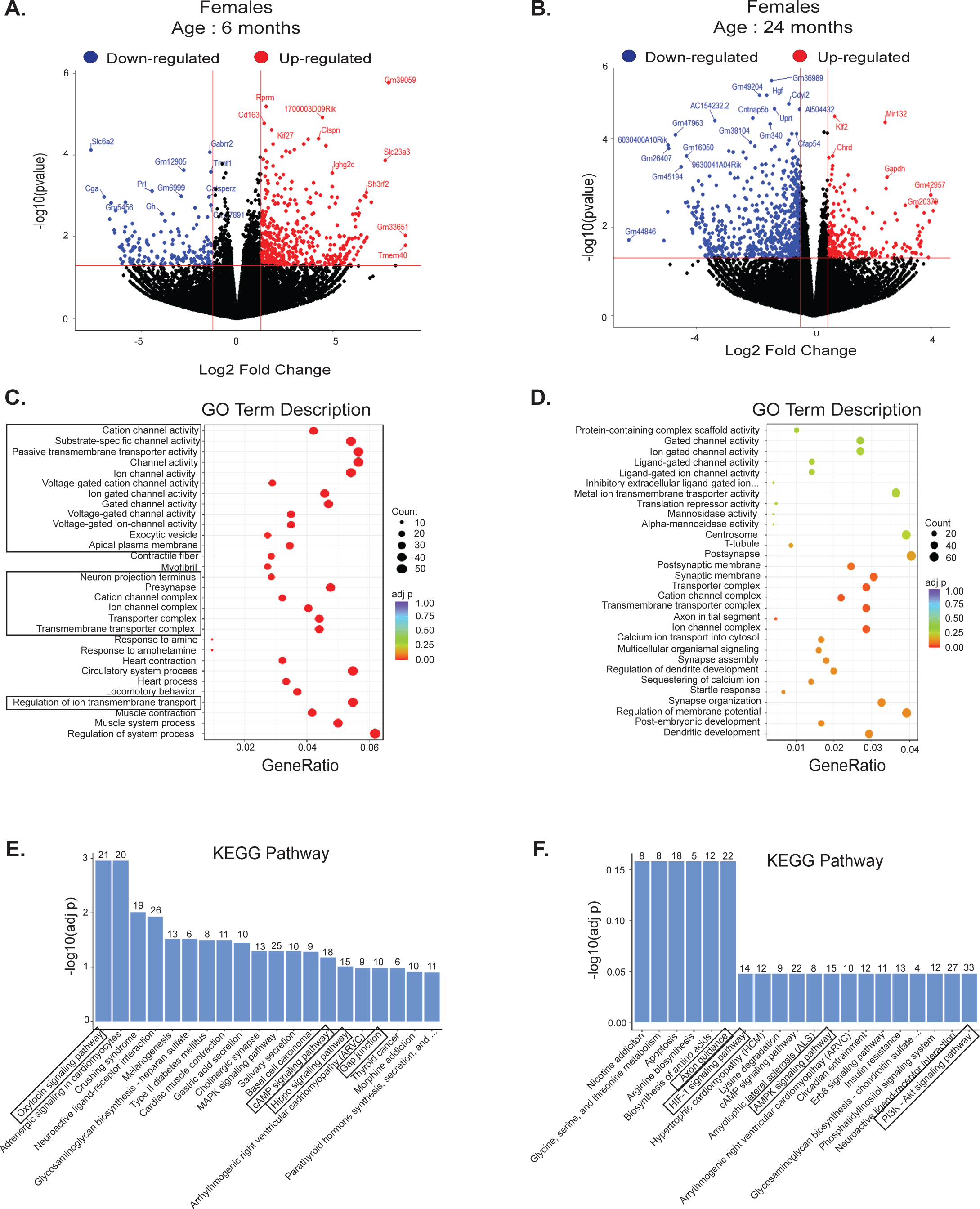
Transcriptome changes in the hippocampi of female cTFEB;HSACre transgenic mice suggest synaptic remodeling and improved synaptic function. (A-B) Volcano plot of differentially expressed genes in hippocampal lysates from female young (A, 6 months old) and aged (B, 24 months old) cTFEB;HSACre mice relative to controls. (C-D) GO term analysis of differentially expressed genes from A and B including gene counts and adjusted p values. Boxes highlight neurotrophic signaling pathways. (E-F) KEGG analysis of differentially expressed genes from A and B. Number of differentially expressed genes (DE) are shown above each category. Boxes highlight categories associated with neural function.

**Figure 7:**
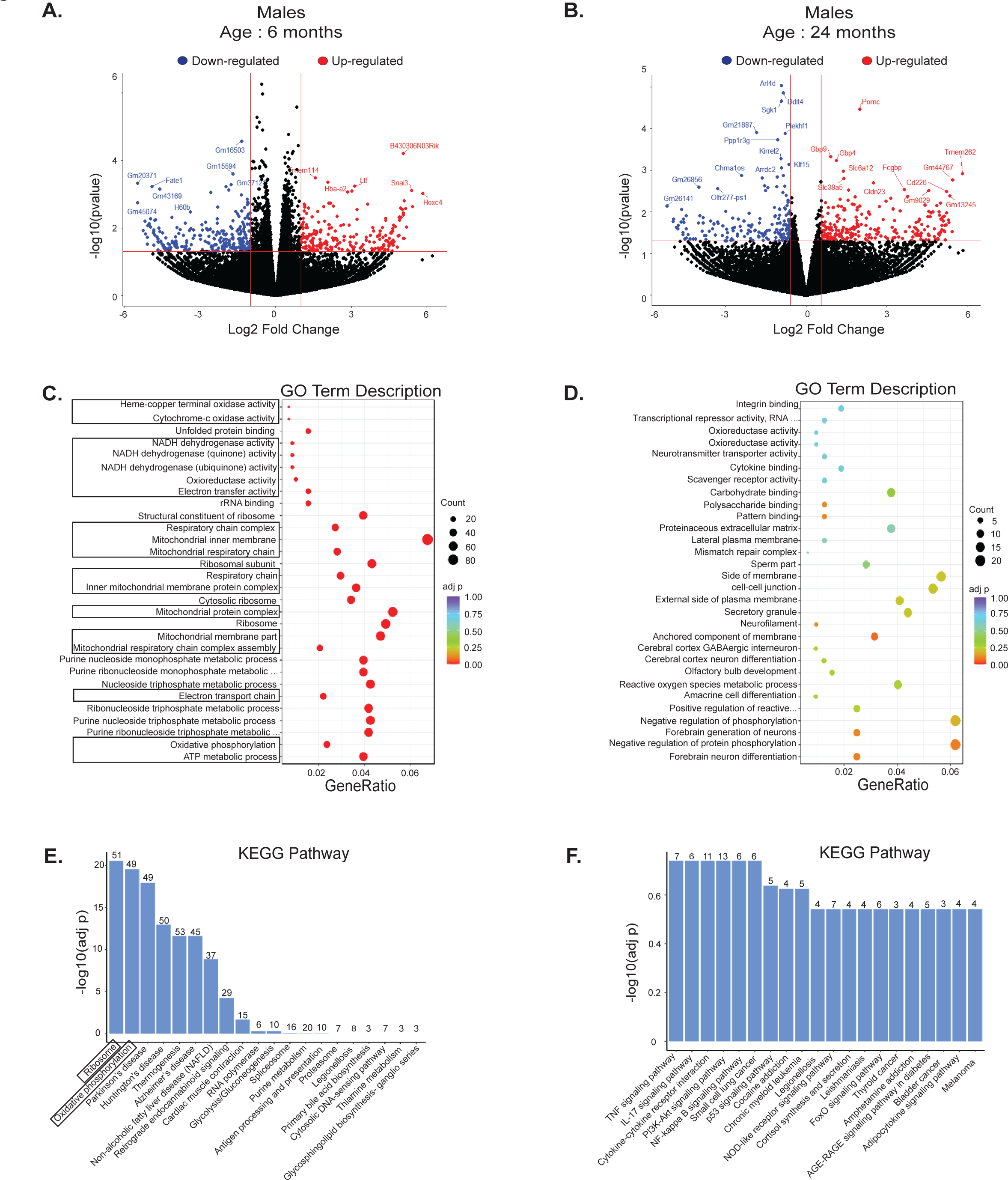
Transcriptome changes in the hippocampi of male cTFEB;HSACre transgenic mice suggest activation of mitochondrial and ribosomal signaling. (A-B) Volcano plot of differentially expressed genes in hippocampal lysates from male young (A, 6 months old, 2 fold change) and aged (B, 24 months old, 1.25 fold change) cTFEB;HSACre mice relative to controls. (C-D) GO term analysis of differentially expressed genes from A and B including gene counts and adjusted p values. Boxes highlight ribosomal and mitochondrial pathways. (E-F) KEGG analysis of differentially expressed genes from A and B. Number of differentially expressed genes (DE) are shown above each category. Boxes highlight ribosomal and mitochondrial pathways.

Bulk RNA-Seq analysis of hippocampal RNA lysates revealed robust changes in gene expression in young cTFEB;HSACre bigenic mice compared to their age- and sex-matched littermate controls (**Figure 6**, **Figure 7, Supplemental Figure 4, and Supplemental Figure 5**). In young female cTFEB;HSACre hippocampi, we detected 618 differentially expressed genes, (DEGs, 435 up-regulated and 165 down-regulated) (**Figure 6A**). Functional GO Enrichment analysis showed these DEGs mapped to key categories associated with synaptic function, including ion- and voltage-gated channel activity, transporter complexes, as well as synapse and synaptic membrane categories (**Figure 6C, boxes**). Enrichment analysis of differential gene expression (KEGG) revealed additional critical categories associated with neural function, including oxytocin signaling, cAMP signaling, Hippo signaling and gap junctions (**Figure 6E**). Furthermore, metabolic and functional pathway analysis (Reactome) also identified key categories known to regulate cognitive plasticity, including GPCR ligand binding, voltage gated potassium channels, and transmission across chemical synapses, among others (**Supplemental Figure 4A**).

Consistent with our biochemical and functional findings suggesting increased neuroprotective benefits in aging cTFEB;HSACre mice (**Figure 5**), we found that these transcriptional changes modulating synaptic function pathways are still present (although not as prominent) in the aging hippocampus of aged female cTFEB;HSACre mice (**Figure 6B, Supplemental Figure 4B&D**). Out of a total of 1765 DEGs, (541 up-regulated and 1224 down-regulated), KEGG functional pathways enriched in our aged female hippocampus included key categories associated with preservation of neuronal activity and cognitive function, including axon guidance, HIF-1 signaling, AMPK signaling and PI3-Akt signaling (**Figure 6F**). We confirmed the age-associated differential expression of a subset of our top-hit genes in a second group of hippocampal RNA lysates using targeted qRT-PCR (**Supplemental Figure 4E**). Altogether, this suggests that overexpression of TFEB in skeletal muscle activates multiple neuroprotective gene networks associated with synaptic function and signaling in the young and aged female hippocampus.

Conversely, in the young male cTFEB;HSACre hippocampus mice we detected 471 DEGs with a 1.25-fold expression change (216 up-regulated and 255 down-regulated), which GO-term analysis associated mostly with multiple mitochondria or ribosomal/transcription targeting pathways (**Figure 7A and C, boxes**). This included mitochondrial respiratory chain, mitochondrial protein complexes, electron transport chain and ATP metabolic processes (**Figure 7C**). Additional enriched metabolic pathways in the hippocampus of young cTFEB;HSACre males included non-sense mediated decay, metabolism of RNA and translation (**Supplemental Figure 5A**). Consistent with this, KEGG analysis revealed the most enriched functional pathways were ribosomes and oxidative phosphorylation (**Figure 7E**). As before, we confirmed the differential expression of a random sampling of our top-hit genes using targeted qRT-PCR (**Supplemental Figure 5E).** When we performed similar analysis on the male aging hippocampus, we detected 539 DEGs in the hippocampus of aged male cTFEB;HSACre mice (278 up-regulated, 261 down-regulated) (**Figure 7B**), with only mild significance on functional pathways (**Figure 7D**, **Figure 7F, Supplemental Figure 5B, and Supplemental Figure 5D**).

Further analysis also revealed that most DEGs appeared to be sex-specific. Out of all DEGs in young cTFEB;HSACre hippocampi (618 in females and 471 in males), there were only 25 shared DEGs, with 16 of these changing in the same direction in both sexes (**Supplemental Figure 6A**). A similar pattern was found in the aged cTFEB;HSACre hippocampus (1765 total DEGs in females and 539 DEGs in males), with 39 shared DEGs and only 3 genes changing in the same direction in both sexes (**Supplemental Figure 6B**). Our results thus strongly suggest a sex bias in the hippocampal transcriptional responses associated with muscle-TFEB overexpression, with the female hippocampi up-regulating neuronal signaling and synaptic function and the male hippocampi upregulating transcriptional and mitochondrial responses. Finally, many of these changes are preserved up until late stages in the lifespan of our model (24 months of age), suggesting that these transcriptional changes may underlie the neurocognitive benefits observed in our aging cTFEB;HSACre cohorts.

## DISCUSSION

Skeletal muscle comprises roughly 40% of the total body mass in a healthy young adult human and is highly susceptible to age-associated metabolic and functional decline ^60,61^. Skeletal muscle health has recently been linked to multiple chronic age-associated conditions, including neurodegenerative diseases ^24,26^, but the role of the so-called muscle-to-brain axis on the pathology of these conditions remains largely unknown. Here, and for the first time in mammals, we provide direct evidence demonstrating the CNS benefits of enhancing muscle-TFEB expression, both during healthy aging and in the context of neurodegenerative proteinopathy.

We found that lifelong overexpression of TFEB, a master regulator of lysosomal and mitochondrial function, reduces age-associated functional and biochemical hallmarks of aging in skeletal muscle ^1^ **(Figure 1-2)**. We confirmed previous reports showing that in young skeletal muscle, acute TFEB-overexpression induces the expression of genes involved in mitochondrial biogenesis, fatty acid oxidation, and oxidative phosphorylation, optimizing mitochondrial substrate utilization and enhancing exercise capacity ^36^. Furthermore, we extend these findings and report that life-long overexpression of muscle-TFEB counteracts age-associated mitochondrial dysfunction in skeletal muscle (**Figure 2**) ^1,44^, ultimately preserving skeletal muscle mass and preventing fiber type switching during aging (**Figure 1**). Indeed, young TFEB-overexpressing muscle had significant increases in the abundance of Type 1 and Type 2A fibers, which are considered to be highly reliant on oxidative phosphorylation and thus more resistant to fatigue. This is consistent with the observed increases in exercise performance of cTFEB;HSACre transgenic mice at both ages (**Figure 1**). Importantly, although we focused our analysis on two key hallmarks of aging (lysosomal and mitochondrial function), our proteomics studies also detected important changes in branched chain amino acid signaling, which have been recently linked to lifespan and healthspan extension in rodents ^62^, indicating that TFEB overexpression may have pan-geroprotective effects on aging skeletal muscle.

The hippocampus is a critical structure in cognitive aging, and hippocampal atrophy and dysfunction are key features of both aging and neurodegenerative diseases ^63,64^. In addition to rescuing age-associated hallmarks in skeletal muscle, we also discovered significant improvements in multiple disease-associated biochemical hallmarks in the hippocampus of cTFEB;HSACre transgenic mice. Indeed, we observed robust benefits on morphological and transcriptional markers of neuroinflammation in the MAPT P301S model of tau toxicity, as well as a significant extension on the lifespan of male MAPT P301S;cTFEB;HSACre transgenic mice (**Figure 4**). We also report important reductions in proteotoxicity and neuroinflammation and improved neurocognitive performance in healthy aging (22-24 month old) cTFEB;HSACre mice (**Figure 5**). Importantly, we observed these benefits without any detectable expression of our transgene in the CNS (**Supplemental Figure 1**). Although previous work examining the muscle-to-brain axis has demonstrated important benefits on the accumulation of misfolded protein inclusions in the CNS ^5,8,19,28^, similar to what we observed in the MAPT P301S;cTFEB;HSACre hippocampus, the profound reductions in neuroinflammation observed in our model suggest modulation of skeletal muscle TFEB-signaling may be a novel target to modulate neuroinflammatory signaling in vivo in the context of aging and/or neurodegenerative diseases.

Previous transcriptional studies of the healthy aging hippocampus have identified dysregulation of ion homeostasis, disruption of neurotransmission, and ribosome biogenesis as key biological process exhibiting differential regulation with aging ^2,65^. Specifically, genes related to synaptic transmission and plasticity appear to be particularly sensitive to age-associated transcriptional dysregulation (reviewed here ^3^). Many of these gene networks are also significantly altered in AD and other associated tauopathies ^57^. It was quite interesting that similar networks were transcriptionally remodeled in the healthy aging cTFEB;HSACre hippocampus, suggesting that these gene networks may be uniquely sensitive to both neurotoxic (aging, neurodegeneration) and neurotrophic (muscle-TFEB overexpression) signaling. Indeed, manipulation of skeletal muscle TFEB modulated expression of similar functional enrichment categories (i.e. trophic factor signaling, neuroinflammation) in the aging and tau-afflicted hippocampus of our cTFEB;HSACre mice in a sex-dependent manner. When we compared KEGG pathways of the differentially expressed genes from the nCounter AD panel in MAPT P301S;cTFEB;HSACre hippocampus with those enriched in the hippocampus of our 24-month old cTFEB;HSAcre mice, we see shared regulation of pathways involved in axon guidance, MAPK signaling, cAMP signaling, and Neuroactive ligand-receptor interactions, suggesting potential converging neurotrophic benefits associated with muscle-TFEB expression.

One of our most exciting (and unexpected) findings is the sex-bias in our CNS and skeletal muscle responses to muscle-TFEB overexpression. In general, we struggled to identify previous literature thoroughly describing female-specific changes to many of these hallmarks of aging on either tissue ^1^. To our surprise, while we were able to validate ‘classical’ age-associated findings such as fiber type switching in aging control males, these phenotypes were less robust or not present in age-matched control females (**Figure 1**), rendering interpretation of the sex-bias of our CNS-targeting benefits difficult. While we ultimately observed neuroprotective effects on both male and female cTFEB;HSACre transgenic mice, it appears that the mechanisms underlying these benefits are not the same. For example, whereas muscle-TFEB overexpression in the MAPT P301S model reduces the accumulation of hyperphosphorylated tau in both sexes, it appears to drive the transcriptional silencing of pro-inflammatory microglial signaling in females, and activate trophic factor signaling in males instead. We also detected similar sex-differences in our healthy aging transcriptional studies, with the female cTFEB;HSACre hippocampus activating pathways associated with synaptic function, while the male cTFEB;HSACre hippocampus activated ribosomal and mitochondrial signaling instead. One potential explanation could be the sex-bias in TFEB expression in skeletal muscle, which is ∼50% higher in male mice, suggesting a threshold effect for local remodeling of skeletal muscle. However, the fact both sexes do display similar exercise endurance at young ages and comparable protection of mitochondrial respiration rates at old ages again suggest converging outcomes of TFEB-mediated geroprotection of aging skeletal muscle. Another consideration is that in our healthy aging studies, both sexes are well into the stage of reproductive senescence, which has been reported to have significant negative impacts on neural function ^66^. While we did not measure circulating levels of sex hormones in our models, there is plenty of evidence for de novo synthesis on neuroactive steroids in the brain having neurotrophic and anti-inflammatory benefits, the phenotypes we also see being modified in our cTFEB;HSACre groups. Interestingly, CNS estrogen appears to control similar signaling pathways as those we detected as differentially expressed in the hippocampus of muscle-TFEB overexpressing female (i.e. synapse function, neurotransmitter signaling) and male (i.e. mitochondrial function) mice ^66^. While it is unclear whether these differential responses between male and female hippocampi reflect local differences in the CNS itself (as recently published here ^67^) or peripheral differences in the expression, secretion or trafficking of circulating signals (such as Ctsb, for example), they underscore the fundamental need to examine sex differences contributing to the biology of aging and neurodegenerative diseases ^68^.

Recent evidence has shown that muscle-originating circulating factors (myokines) appear to play central roles in regulating CNS health and function. Ctsb is secreted from skeletal muscle into circulation in response to exercise, and is required for the full manifestation of exercise-associated benefits on the CNS, including increases in hippocampal Bdnf and the activation of neurogenesis ^20^. Circulating Ctsb has also been proposed to cross the blood-brain barrier ^20^, and appears to remodel the extracellular matrix ^69^, enhancing axonal outgrowth through degradation of chondroitin sulfate ^70,71^. Furthermore, high Ctsb expression levels in the hippocampus have been reported in low-anxiety mouse lines ^72,73^, suggesting that Ctsb may play important roles in maintaining neuronal homeostasis in brain regions with high relevance for both aging and age-associated neurodegenerative disease. Through our proteomics in silico analysis, we identified Ctsb as our highest expressed secreted factor in cTFEB;HSACre skeletal muscle, and we confirmed higher levels of circulating Ctsb in the serum of cTFEB;HSACre transgenic mice. Our results imply that additional research into the role and function of circulating proteases and their ability to remodel the CNS may be warranted. Interestingly, recent evidence demonstrates pronounced involvement of the (macro)autophagy molecular machinery in cellular secretion ^74^. Indeed, key signaling molecules and cytokines associated with inflammation, such as IL1B (interleukin 1 beta) and IL6 (interleukin 6), are released into the extracellular space via autophagy-dependent secretion. There is also a growing body of evidence that age ^46^ and exercise ^75^, known modifiers of skeletal muscle metabolism, alter the composition of the skeletal muscle secretome. This suggests that activation of TFEB-overexpression may be modifying the skeletal muscle secretome, potentially enriching it with neurotrophic factors or depleting it from neurotoxic factors, although this hypothesis remains to be tested.

Over the last ten years, there has been growing evidence that suggests prominent contributions of the periphery to the etiology of neurodegenerative diseases. Our discovery that skeletal muscle TFEB-signaling can be a crucial site for regulation of CNS health and function provides compelling evidence for a new therapeutic delivery avenue into the brain, providing a currently unexplored new diagnostic and therapeutic intervention site (i.e. skeletal muscle) for preserving brain health.

## Authorship Contributions

IM, CJC, AG, HM, AB, AH, KL, EL, KC, MM, DP, MB, AN, NN, GM, LW, MM, NM, EB and EF: Performed the experimental work. Reviewed and edited the manuscript.

CJC and IM: Wrote and edited the manuscript.

CJC and ALS: Conceptualized the experiments outlined here and obtained all associated funding.

## Supplemental Figures

**Supplemental Figure 1: Skeletal muscle-restricted expression of human TFEB protein.**

(A-B) qRT-PCR analysis for human *TFEB* mRNA expression in 6-month old (A) and 21-24 month old (B) control and cTFEB;HSACre quadriceps skeletal muscle, heart ventricles and hippocampus.

(C) Representative images of gastrocnemius cross sections stained for Hoechst (blue) and Human TFEB (white). Insets of nuclear hTFEB staining are 5x zooms of the areas denoted by the yellow squares. Scale bars = 100 µm.

(D-G) Immunoblot analysis for 3x-FLAG TFEB protein expression from lysates of brown adipose (D), liver (E), heart ventricles (F), and brain (G) tissue from 6-month old female cTFEB;HSACre mice and control littermates. cTFEB;HSACre skeletal muscle lysates are shown in the first lane of each blot as a positive control for 3x-FLAG TFEB expression. FLAG-eGFP protein cassette (arrow) as well as non-specific bands (asterisks) are indicated.

(H) Representative images of quadriceps and gastrocnemius cross sections from 7-week old male TdTomato;HSACre transgenic mice stained for Hoechst (blue), TdTomato (red), and Laminin (white). Scale bars = 100 µm.

(I) Representative images of hemibrain sections from 7-week old male TdTomato;HSACre transgenic mice stained for Hoechst (blue), TdTomato (red). Insets of hippocampal regions are 5x zooms of the areas denoted by the yellow squares. Scale bars = 1 cm.

Controls are age-matched littermates. Each point represents the average of all data collected from an individual. Data are represented as mean ± SEM. One-way ANOVA, post-hoc multiple comparisons * p<0.05, ** p<0.01, *** p<0.001. Lack of annotation indicates comparisons were not significant.

**Supplemental Figure 2: Fiber typing metrics in cTFEB;HSACre transgenic muscle.**

(A-H) Feret’s minimum diameter (A-D) and fiber cross sectional area (E-H) of all 4 fiber types in gastrocnemius muscle of young (7-9 months) and aged (22-24 months) control and cTFEB;HSACre transgenic mice of both sexes.

(I) Immunoblot analysis for LC3 and total ubiquitin in quadriceps protein lysates from young (5-6 months) and aged (22-24 months) control and cTFEB;HSACre mice of both sexes. Total Protein Stain Free lanes are shown as a loading control. Marker densitometry quantification relative to Protein Stain Free densitometry shown below.

Controls are age-matched littermates. Each point represents the average of all data collected from an individual. Data are presented as mean ± SEM. * p<0.05, ** p<0.01, **** p<0.0001 ANOVA and interactions of sex x age post-hoc. Lack of annotation indicates comparisons were not significant.

**Supplemental Figure 3: Muscle-TFEB overexpression reduces hyperphosphorylated tau accumulation and modifies expression of AD-associated genes in the hippocampus of MAPT P301S transgenic mice.**

(A) Representative images of the hippocampal dentate gyrus from 9-month old control (top), MAPT P301S (middle), and MAPT P301S;cTFEB;HSACre (bottom) mice stained for Hoechst (blue) and AT180 PhosphoTau (white). Insets of hippocampal regions are 5x zooms of the areas denoted by the yellow squares. Representative images are from males. Scale bars= 100 µm.

(B-C) Volcano plot of differentially expressed genes in the hippocampi of 9-month old male (B) and female (C) MAPT P301S;cTFEB:HSACre animals relative to age-matched P301S MAPT littermates using the Nanostring nCounter AD panel. Down-regulated genes in blue, up-regulated genes in red.

(D) Cumulative ataxia score of 9-month old control, MAPT P301S, and MAPT P301S;cTFEB;HSACre mice of both sexes.

(E) Weight (in grams) of 11-month old control MAPT P301S, and MAPT P301S;cTFEB;HSACre mice of both sexes.

Controls are age-matched littermates. Each point represents the average of all data collected from an individual. Data are represented as mean ± SEM. One-way ANOVA, post-hoc multiple comparisons * p<0.05, ** p<0.01. Lack of annotation indicates comparisons were not significant.

**Supplemental Figure 4: Functional analysis of differentially expressed genes in the hippocampus of female cTFEB;HSACre transgenic mice.**

(A-B) Reactome analysis of differentially expressed genes in hippocampal mRNA lysates from female young (6 months old, A) and aged (24 months old, B) cTFEB;HSACre mice. Number of differentially expressed genes (DE) are shown above each category.

(C-D) GO enrichment analysis of differentially expressed genes from A-B. Number of differentially expressed genes (DE) are shown above each category.

(E) qRT-PCR analysis for select DE genes in hippocampal mRNA lysates from female young (6 months old, solid circles) and aged (24 months old, open circles) control (gold) and cTFEB;HSACre (purple) mice. Expression levels normalized to young controls.

Controls are age-matched littermates. Each point represents the average of all data collected from an individual. Data are represented as mean ± SEM, O One-way ANOVA, post-hoc multiple comparisons, * p<0.05, ** p<0.01. Lack of annotation indicates comparisons were not significant.

**Supplemental Figure 5: Functional analysis of differentially expressed genes in the hippocampus of male cTFEB;HSACre transgenic mice**

(A-B) Reactome analysis of differentially expressed genes in hippocampal mRNA lysates from male young (6 months old, A) and aged (24 months old, B) cTFEB;HSACre mice. Number of differentially expressed genes (DE) are shown above each category.

(C-D) GO enrichment analysis of differentially expressed genes from A-B. Number of differentially expressed genes (DE) are shown above each category.

(E) qRT-PCR analysis for select DE genes in hippocampal mRNA lysates from female young (6 months old, solid circles) and aged (24 months old, open circles) control (gold) and cTFEB;HSACre (purple) mice. Expression levels normalized to young controls.

Controls are age-matched littermates. Each point represents the average of all data collected from an individual. Data are represented as mean ± SEM, One-way ANOVA, post-hoc multiple comparisons, * p<0.05, ** p<0.01. Lack of annotation indicates comparisons were not significant.

**Supplemental Figure 6: Overlap of differentially expressed genes confirms sex-specific transcriptional remodeling in the hippocampus of TFEB;HSACre transgenic mice.**

(A) Overlap of differentially expressed genes (DEGs) in hippocampal mRNA lysates from young (6 months old) cTFEB;HSACre mice relative to their age-matched controls. Number of up-or down-regulated DEGs are shown next to each unique and shared category.

(B) Overlap of differentially expressed genes (DEGs) in hippocampal mRNA lysates from aged (24 months old) cTFEB;HSACre mice relative to their age-matched controls. Number of up-or down-regulated DEGs are shown next to each unique and shared category.

## STAR*Methods

### Key resources table

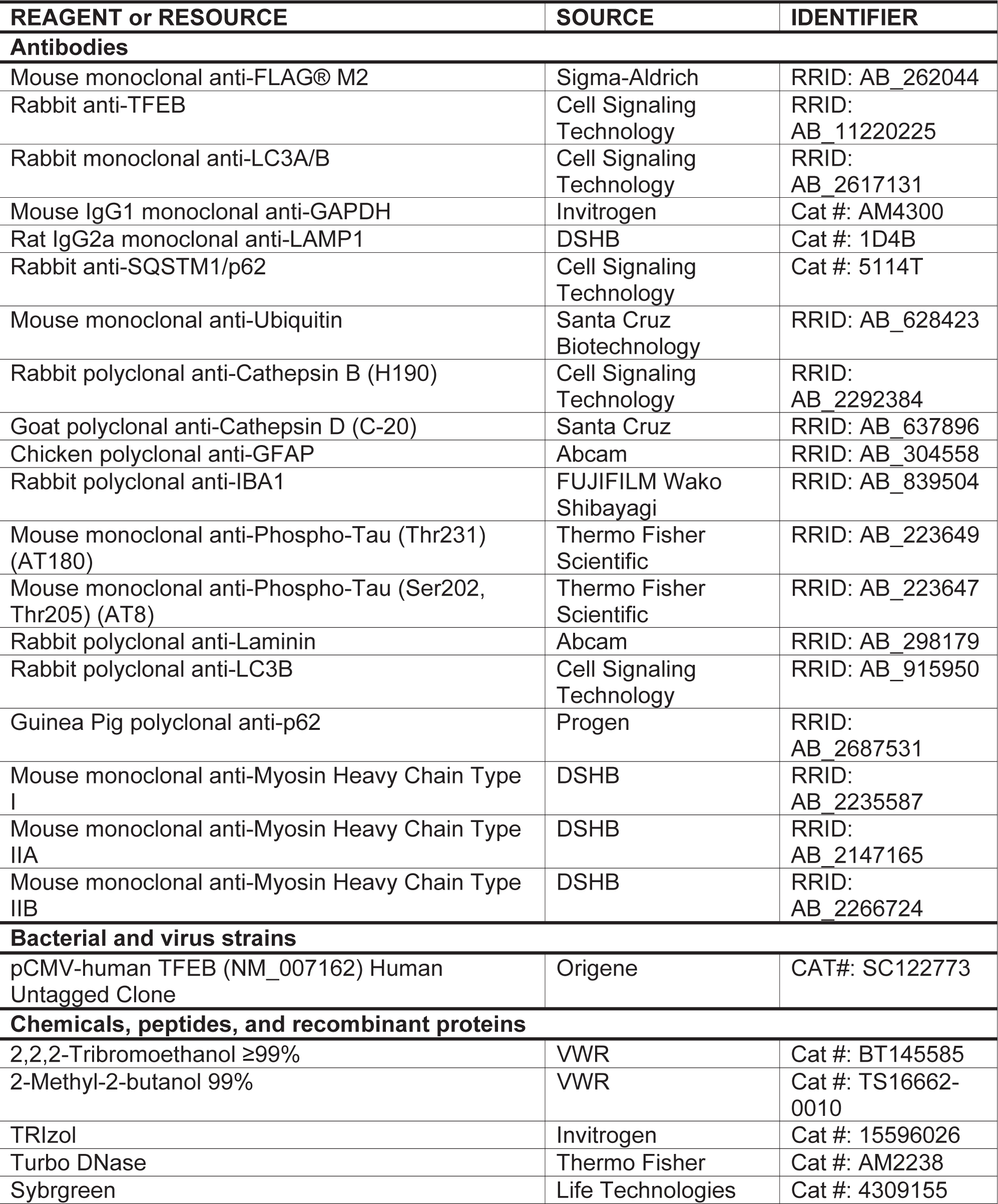
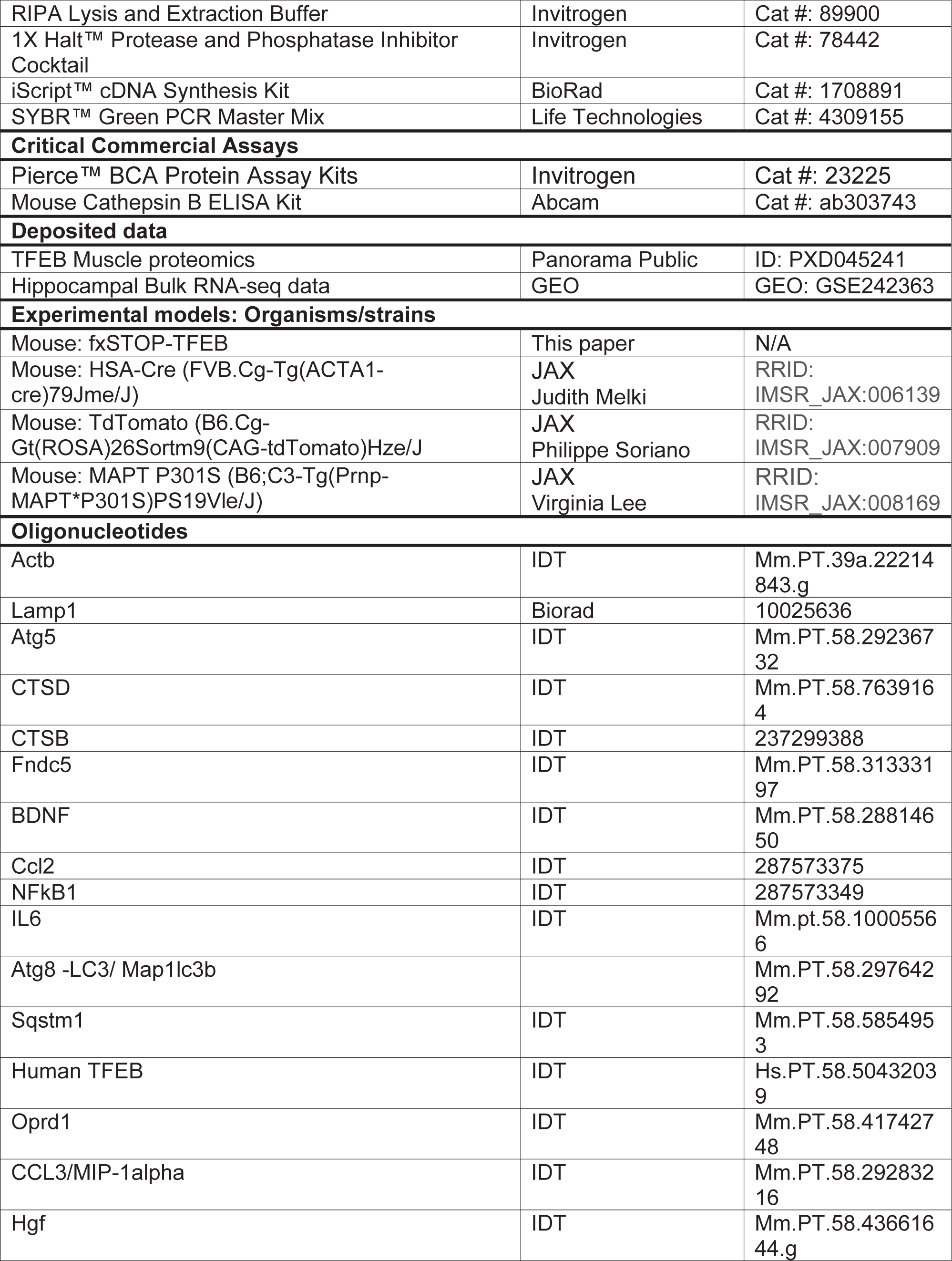
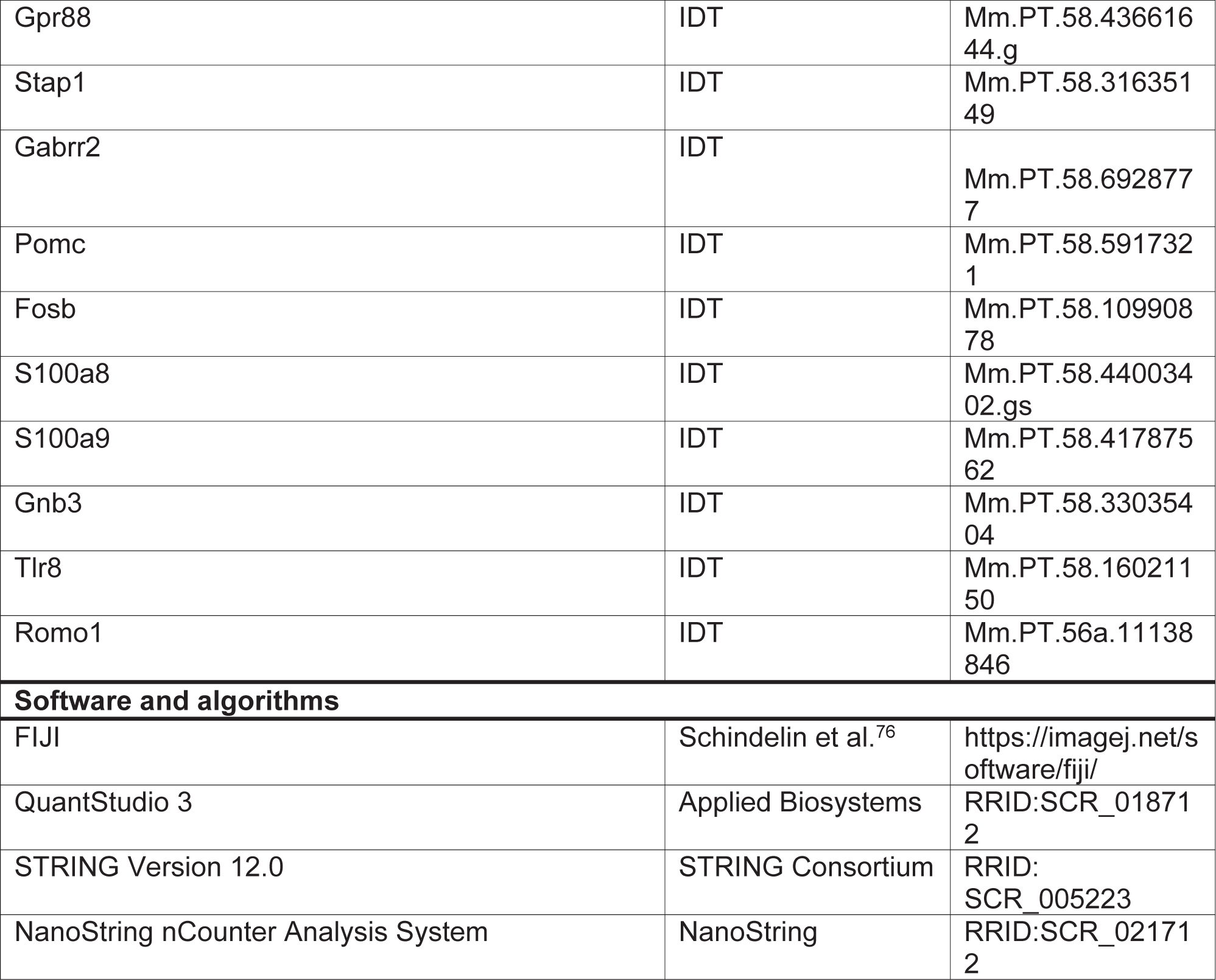

### Resource availability

#### Lead contact

Further information and requests for resources and reagents should be directed to and will be fulfilled by the lead contact, Constanza Cortes.

### Materials availability

The fxSTOp-TFEB line of transgenic mice is available for sharing with academic institutions free-of-charge. For-profit institutions will adhere to UCSD’s policy regarding copyrighted material.

## Data and code availability

- Bulk RNA-seq data have been deposited at GEO and are publicly available as of the date of publication. Accession numbers are listed in the key resources table. Muscle proteomics data have been deposited at Panorama Public and are publicly available as of the date of publication. ID numbers are listed in the key resources table. Microscopy data reported in this paper will be shared by the lead contact upon request.
- This paper does not report original code.
- Any additional information required to reanalyze the data reported in this work paper is available from the lead contact upon request.

### Experimental model details

Mouse models used were the novel fxSTOP-TFEB transgenic line generated for this publication, the HSA-Cre transgenic mice (JAX Strain No: 006139 | HSA-Cre79) obtained from JAX, TdTomato mice (B6.Cg-Gt(ROSA)26Sortm9(CAG-tdTomato)Hze/J, JAX Strain #:007909) also from JAX, and the MAPT P301S line (Stock No: 008169) which originated in the laboratory of Dr. Virginia Lee. All animals have the C57BL/6J genetic background, both males and females were used wherever possible.

### Method details

#### Generation of fxSTOP-TFEB transgenic mice

We used a pCMV-human TFEB expressing vector from Origene (clone # sc122773), used previously ^27^. We cloned a 3x-FLAG fragment via insertion into the Acc65I and Pac1 restriction sites of the targeting vector. The fxSTOP-TFEB vector was generated in multiple steps, as follows: (1) 5’ and 3’ targeting arms were inserted by PCR onto the ends of a minimal 1.8 kb, chloramphenicol-resistant vector; (2) an ampicillin resistance cassette was cloned into the MluI, XmaI sites of the targeting vector; (3) an 1.1 kb fragment encoding the actin promoter, a loxP site, a 3x-FLAG tag fused to EGFP was cloned into the NheI, PacI restriction sites of the targeting vector; (4) a 2.8 kb fragment containing the second loxP site, followed by 3x-FLAG-human TFEB sequence. The final fxSTOP-FLAG-human TFEB vector was microinjected into C57BL/6J:C3H/HeJ F1 hybrid oocytes. Of six founders identified, two gave rise to fxSTOP-FLAG-human TFEB lines with comparable phenotypes. Of these lines, we chose to focus our studies principally on line 419. The presence of the floxed 3x-FLAG-eGFP STOP-3x-FLAG-human TFEB cassette is verified by allele dependent qPCR genotyping analysis with each generation (Transnetyx), and expression of the 3x-FLAG-eGFP sequence (upstream of the STOP codon) in single transgenic fxSTOP-TFEB mice is assessed every other generation. We confirmed excision of 3x-FLAG-EGFP-STOP cassette in the presence of Cre-recombinase by crossing fxSTOP-TFEB transgenic female mice with HSA-Cre transgenic mice (JAX Strain No: 006139 | HSA-Cre79), and corresponds to the HSA-Cre driver mouse line from the laboratory of Dr. Judith Melki. TdTomato mice (B6.Cg-Gt(ROSA)26Sortm9(CAG-tdTomato)Hze/J, JAX Strain #:007909) express robust tdTomato fluorescence following Cre-mediated recombination, is congenic on the C57BL/6J genetic background and were also obtained from JAX. The MAPT P301S line (Stock No: 008169) originated in the laboratory of Dr. Virginia Lee. cTFEB;HSACre;MAPT P301S transgenic mice were generated by crossing fxSTOP-TFEB females to MAPT P301S males. Double transgenic offspring was then crossed with heterozygote HSA-Cre transgenic individuals. All commercially available lines were obtained directly from JAX. All lines have been maintained in a C57/B6 background for over ten generations in our lab. Mice were housed in a temperature-and humidity-controlled environment on a 12-hour light/dark cycle with food and water ad libitum. Mice of both sexes were used for all experiments in equal numbers, or as close to equal numbers as possible. For all MAPT P301 experiments, control animals are MAPT P301S;fxSTOP-TFEB-;HSACre- and/or MAPT P301S;fxSTOP-TFEB+;HSACre-littermates. For all aging experiments, control animals are fxSTOP-TFEB-;HSACre- and/or fxSTOP-TFEB+;HSACre-littermates. In our hands, we did not detect any differences on tau or aging-associated CNS phenotypes on fxSTOP-TFEB+ animals compared to the wild-type controls. All animal experimentation adhered to NIH guidelines and was approved by and performed in accordance with the University of California, San Diego, Duke University, University of Alabama at Birmingham, and the University of Southern California Institutional Animal Care and Use.

#### Tissue Collections

Animals were anesthetized with 3.8% Avertin Solution. All animals received a transcardial perfusion with 60 mLs of cold 1x PBS. Half of CNS tissue (right/left hemi-brain) was post-fixed in 4% PFA for less than 24 hours before being transferred to 30% sucrose for another 24 hours before cryo-sectioning and staining the other half was micro-dissected into cortex and hippocampus and flash-frozen in liquid nitrogen for RNA and protein analyses. Half of skeletal muscle tissue (right/left limb muscle) was fresh frozen in OCT following dissection, while the other half was flash-frozen in liquid nitrogen for RNA and protein analyses.

#### RT-PCR Analysis

Flash frozen perfused isolated mouse tissues (quadriceps, hippocampus) were placed in plastic tubes with silica beads (MP Biomedical, 116913100) and homogenized in the FastPrep-24^TM^ 5G bead beating grinder and lysis system (MP Biomedical, 116005500) in TRIzol (Invitrogen, 15596026). RNA was extracted via standard phenol-chloroform procedures followed by DNase digestion (ThermoFisher, AM2238). cDNA was generated using iScript (Bio-Rad Technologies, 1708891). Quantitative PCR was run using Sybrgreen (Life Technologies, 4309155) for chemical detection (Applied Biosystems; QuantStudio 3). Enrichment was determined based on double-delta CT value. Primers were ordered from IDT Predesigned qPCR Assays unless otherwise specified. Primers used are listed in the Key Resources Table.

#### Protein Extraction and ImmunoBlot Analysis

Protein lysates from muscle tissue was prepared as previously described ^27,28^. In short, Flash frozen perfused isolated mouse tissues (quadriceps, hippocampus) were placed in plastic tubes with silica beads (MP Biomedical, 116913100) and homogenized in the FastPrep-24^TM^ 5G bead grinder and lysis system (MP Biomedical, 116005500) in RIPA Lysis and Extraction Buffer (Invitrogen, 89900), 1X Halt^TM^ Protease and Phosphatase Inhibitor Cocktail (Invitrogen, 78442), and 1% SDS. Protein concentration was quantified using a Pierce™ BCA Protein Assay (23227). For brain tissue, Fifty or 25 µg of homogenized proteins were loaded per lane, and after running Any KD, 10%, or 4-15% Mini-PROTEAN TGX Gels (BioRad, 4568124, 4561034, and 4561084), samples were transferred to 0.45 µm PVDF membranes (BioRad, 1704275), which were blocked in 5% BSA in PBS at RT for 1 hr. For skeletal muscle, proteins were separated on 4-20% Stain-Free TGX gels (Bio-Rad, 5678092), separated to PVDF, and blocked in 5% non-fat dry milk in TBST (1X TBS with 0.05% Tween-20) for 1 hour at RT. Membranes were incubated with anti-FLAG antibody (Sigma, M2, F1804, 1:1000), anti-TFEB antibody (Cell Signaling, 4240,1:1000), anti-Lamp1 antibody (DSHB, 1DB4, 1:1000), anti-LC3A/B antibody (Cell Signaling, 12741, 1:1000), anti-P62 antibody (Cell signaling, 5114T, 1:1000), anti-ubiquitin antibody (Santa Cruz, sc-8017, 1:1000), anti-Cathepsin B antibody (Cell signaling, 3383S, 1:1000), anti-Cathepsin D antibody (Santa Cruz, sc-6486, 1:1000),or anti-GAPDH (Invitrogen, AM4300, 1:5000) in PBS-T with 5% BSA at 4°C overnight. The primary antibody was visualized with horseradish-peroxidase conjugated anti-rabbit at (Cell Signaling, 7074P2, 1:5,000) and enhanced chemiluminescence (BioRad, 170-5060) or goat-anti-mouse IgG 680 (Invitrogen, A21058, 1:10,000). Densitometry analysis was performed using the BioRad Image Lab 6.1 software application.

#### Immunofluorescence Studies

Tissue was embedded in OCT (TissueTek, 4583), frozen utilizing 2-methylbutane (VWR, 103525-278) submerged in liquid nitrogen and stored at −80 C until used. All samples were sectioned on a standing cryostat (Leica). Brain sections were 20 microns thick, while muscle sections were 15 microns thick. For immunohistochemistry, sections were permeabilized with .25% Triton for 15 mins and blocked with 4% BSA for 30 minutes-1 hour. Brain sections were then incubated with anti-GFAP (ab4674, Abcam, 1:200), anti-IBA1 (Wako, 019-19741, 1:200), anti-AT180 TAU (ThermoFisher, MN1040, 1:100) and anti-AT8 TAU (ThermoFisher, MN1020, 1:100) while muscle sections were incubated with anti-laminin ( Abcam, ab11575, 1:200), anti-FLAG (Sigma, F1804, 1:1000), anti-LC3B (Cell Signaling, 2775S, 1:200), anti-LAMP1 (Novus, NB100-77685, 1:200), anti-p62 (Progen, GP62-C, 1:200), anti-MHC Type 1 (DSHB, BD-D5, 1:100), anti-MHC Type 2A (DSHB, SC-71, 1:100), anti-MHC Type 2B (DSHB, BF-F3, 1:100), and anti-hTFFEB (CST, 4240, 1:200) overnight at 4°C and incubated with secondary antibodies at RT for 1 hour (both diluted in 4% BSA). Next the slides were washed with Hoescht (ThermoFisher, 62249, 1:5000) and mounted with prolong glass (Invitrogen, P36984). All slides were washed with 1X PBS three times for 5 minutes each between steps.

Slides with brain sections stained for TAU and glia markers were imaged in the UAB HIRF HRIF Confocal/Light Microscopy Core utilizing the 10x objective on the Nikon A15R/SIM Confocal microscope. Z-stacks of the entire hippocampus area/section were collected, and max intensity projections were generated using FIJI. Lipofuscin imaging was performed using an epifluorescent Nikon light microscope with a 20x objective. For TdTomato imaging, stitched whole-section z-stacks were acquired using a Nikon A1R HD25 confocal microscope of nuclei in the blue channel and native TdTomato fluorescence in the red channel. Muscle sections were imaged with an ECHO Revolution epifluorescent microscope.

For quantification of astrocyte and/or microglia parameters, NIS elements was used to collect morphometric information and numbers. A size exclusion parameter was used for GFAP positive objects under 10 microns, IBA-1 positive objects under 15 microns, and any object larger than 5,000 microns were all excluded due to standard assessments of cell sizes. FIJI was used for quantification of lipofuscin and LAMP1 puncta. Muscle fibertype composition, number, and size were quantified using Myosoft ^77^, a FIJI plugin.

### Quantitative Proteomics Sample Preparation and Data-independent Acquisition Mass Spectrometry

#### Lysis/Digestion

We identified distinct protein species from crude total extractions of whole quadricep muscle by mass spectrometry. Quadricep muscle powder lysate in 50 mM ammonium bicarbonate with 0.1% RapiGest (Waters, Cat# 186001861) were processed and digested using Qiagen Tissue Lyser II (Qiagen) following the manufacturer’s protocol. Protein concentrations were determined using Pierce BCA Protein Assay Kit (ThermoFisher, PI23227). 150 μg of lysate was reduced with DTT (Sigma, Cat # D0632), alkylated with IAA (Sigma, Cat # D0632) and digested at 1 μg trypsin (Thermo, Cat # PI-90057) to 50 μg protein ratio for 16 hours at 37°C. Digests were acidified with 200 mM HCl (Fisher, Cat # A144), cleaned with MCX columns (Waters, Cat# 186000782), dried with a vacuum concentrator, and reconstituted in 0.1% formic acid (Fisher, Cat# A117) in water.

#### Liquid Chromatography and Mass Spectrometry

One μg of each sample with 150 fmol of Pierce Retention Time Calibrant (PRTC, Thermo, Cat # PI-88321) was loaded onto a 30 cm fused silica picofrit 75 μm column (New Objective, Woburn, MA, Cat # PF3607510N5) and 3.5 cm 150 μm fused silica Kasil1 (PQ Corporation) frit trap loaded with 3 μm Reprosil-Pur C18 (ESI Source Solutions, Cat # r13.aq.0001) reverse-phase resin analyzed with a Thermo Easy nano-LC 1000 coupled to a Thermo Orbitrap Fusion Lumos Mass Spectrometer. Data-independent (DIA) mass spectrometry with gas-phase fractionation was performed as described previously ^78,79^.

#### Data Analysis

Thermo DIA MS/MS RAW files were converted to mzML format using Proteowizard (version 3.0.19045) ^80^ using vendor peak picking and demultiplexing. Chromatogram spectral libraries were created using default settings (10 ppm tolerances, trypsin digestion, HCD b- and y-ions) of Walnut in EncyclopeDIA (version 0.6.14) ^81^ using the Uniprot mouse canonical FASTA ^82^. Quantitative spectral libraries were created by mapping spectral to the chromatogram spectral library using EncyclopeDIA requiring a minimum of 3 quantitative ions and filtering peptides at a 1% FDR using Percolator 3.01 ^83^. The quantitative spectral library is imported into Skyline-daily 4.2.1.19058 ^84^ with the Uniprot mouse canonical FASTA as the background proteome to map peptides to proteins. The mzML data is imported and all data is TIC normalized.

#### Data and code availability

The Skyline document and raw files for the proteomics DIA MS/MS data are available at Panorama Public ^85^. ProteomeXchange ID: PXD045241. Access URL: https://panoramaweb.org/mouse-muscle-TFEB-proteomics.url. All original data has been deposited at Panorama and is publicly available as of the date of publication. Any additional information required to reanalyze the data reported in this paper is available from the lead contact upon request.

### High-Resolution Respirometry and ROS Production

High-resolution respirometry (HRR) was conducted on a G-model Oxygraph-2k machine (Oroboros Instruments, Innsbruck, Austria). Reagents, muscle fiber preparation, and Substrate Uncoupler Inhibitor Titration (SUIT) protocols used for HRR have been previously described ^86^. Antimycin A was sufficient to block complex 3 activity; therefore, myxothiazol was not added in our SUIT protocol. ROS production was detected using the O2k-Fluo (Oroboros Instruments) and a modified Amplex Red assay. After daily background correction, chambers were closed and 20 μM final concentration Amplex Red ultra (Life Technologies, A36006), 1 U/mL horseradish peroxidase (Sigma-P8250), and 5 U/mL superoxide dismutase (Sigma-S8409) were added to the wells. After the amp-slope stabilized 0.1 μM H2O2 (Sigma-95321) was added for calibration. Throughout the entire assay, repeated injections of 0.1 μM H2O2 were used for calibration including one injection after the muscle tissue was added to correct for autofluorescence. All subsequent 0.1 μM H2O2 injections were used to control for changing concentrations of Amplex Red during the assay. Samples for HRR aere conducted in duplicate and means were used for the final analyses. To eliminate confounds from tissue damage or hyperpermeabilization, only samples with stable oxygen flux and Raw-amps were included in the final analysis.

### Gene Expression Analysis

#### RNA Sequencing and Bioinformatics Analysis

RNA was extracted as mentioned above from flash frozen hippocampal tissue for four biological replicates (sex/genotype) and checked for quality by bioAnalyzer by Novogene (Novogene Corporation INC, Sacramento, CA, United States). mRNA libraries were prepared by Novogene using poly-T oligo-attached magnetic beads. cDNA was synthesized with random hexamer primers and dUTP or dTTP depending on library directionality. For the both non-directional and directional library prep; end repair, A-tailing, adapter ligation, size selection, amplification, and purification were performed. For the directional library, end repair, USER enzyme digestion was also performed. Libraries were checked via Qubit and qPCR for quantity and by bioanalyzer for size. Quantified libraries were pooled and bulk RNA-seq analysis was performed via NovaSeq PE150 (Illumina, San Diego, CA, United States) high throughput sequencing. Bioinformatics analysis for differentially expressed genes was performed by Novogene. Raw reads were cleaned to remove adapters, poly-N and low qualities reads from the data. The reference index was built using Hisat2 v2.05 and pair-end cleaned reads were aligned to the reference genome GRCm39 through Hisat2 v2.05. reads were counted using featureCounts v1.5.0-p3. FPKM of each gene was calculated based on the length of the gene and read counts mapped. Differential expression analysis was performed using the DEseq2 R package (1.20.0). Gene Ontology (GO) and KEGG enrichment analysis of differentially expressed genes was implemented by the clusterProfiler R package. For all analysis methods and graphic representations, a p-value less than 0.05 was considered significantly enriched by differentially expressed genes. For young (6-month) animals, a 2-fold change was used to determine differentially expressed genes. A 1.2 fold change was used for aged (24-month) animals. Volcano plots were generated in RStudio using ggplot2 using these criteria.

#### Nanostring nCounter AD panel

Assays were performed with 100 ng aliquots of RNN, extracted using the PureLink RNA extraction kit (Invitrogen) and run via the NanoString nCounter Analysis system (NanoString Technologies, Seattle, WA, USA) at the UAB Nanostring Core, following previously described and established protocols^54^. After codeset hybridization overnight, the samples were washed and immobilized to a cartridge using the NanoString nCounter Prep Station. Cartridges were scanned in the nCounter Digital Analyzer at 555 fields of view for the maximum level of sensitivity. Raw reads and the Nanostring Mm_AD30_v1.1 probe annotation codeset were imported into the Nanostring proprietary software, nSolver 4.0, and QC was performed following company protocol guidelines. Counts for target genes were normalized to house-keeping genes included in the panel (Cltc, Gapdh, Gusb, Hprt, Pgk1, and Tubb5). Specifically, background correction was performed using the negative control at the cutoff of mean + 2 standard deviation. Housekeeping genes were used for normalization based on geometric mean, Gene expression was normalized using NanoStringNorm R package. All p-values were adjusted using a false discovery rate (FDR) correction of 1% for multiple comparisons. Differential expression and pathway scoring analysis, as well as graphical QC representation were performed in the nSolver Advanced Analysis Software 2.0 using a custom analysis to utilize generated normalized counts. Differential gene expression values were presented asMAPT P301S baseline compared to the MAPT P301S;cTFEB;HSACre cohort for each sex, and a p value of 0.05 or less was described as significant. Volcano plots were generated with ggplot2 in Rstudio using significant genes and a cutoff of 1.2 or greater fold change. The pathway scoring module summarizes a single score from the data from a pathway’s genes ^87^.

#### Data and code availability

Bulk RNA-seq data have been deposited at GEO and are publicly available as of the date of publication. Accession numbers (GSE242363) are listed in the key resources table. No new code was generated or used in the analysis of this dataset. Any additional information required to reanalyze the data reported in this paper is available from the lead contact upon request.

#### Cathepsin B ELISA Analysis

Fifty μl of serum (collected via blood cardiac puncture and isolated via standard coagulation/centrifugation protocols) was diluted 1:1 and processed with the Cathepsin B ELISA kit (Abcam, ab119585). Diluted samples were processed following kit instructions, and imaged at 450 nm (Tecan, Infinite M plex). Amount of cathepsin B was determined using a standard dilution curve of known concentration.

### Mouse Phenotyping and Behavioral Studies

#### Barnes Maze

The Barnes maze apparatus is an opaque Plexiglas disc 75 cm in diameter elevated 58 cm above the floor by a tripod. Twenty holes, 5 cm in diameter, are located 5 cm from the perimeter, and a black Plexiglas escape box (19 x 8 x 7 cm) is placed under one of the holes. Distinct spatial cues are located all around the maze and are kept constant throughout the study. On the first day of testing, a training session was performed, which consists of placing the mouse in the escape box for one minute. After the one-minute habituation period, the first session was started. At the beginning of each session, the mouse was placed in the middle of the maze in a 10 cm high cylindrical black start chamber. After 10 seconds the start chamber was removed, a buzzer (80 dB) and a light (400 lux) were turned on, and the mouse was set free to explore the maze. The session ended when the mouse entered the escape tunnel or after 3 min elapsed. When the mouse entered the escape tunnel, the buzzer was turned off and the mouse was allowed to remain in the dark for one minute. When the mouse did not enter the tunnel by itself it was gently put in the escape box for one minute. The tunnel was always located underneath the same hole (stable within the spatial environment), which was randomly determined for each mouse. Mice were tested once a day for 4 days for the acquisition portion of the study. For the 5th test (probe test), the escape tunnel was removed and the mouse was allowed to freely explore the maze for 3 min. The time spent in each quadrant was determined and the percent time spent in the target quadrant (the one originally containing the escape box) was compared with the average percent time in the other three quadrants. This was a direct test of spatial memory as there was no potential for local cues to be used in the mouse’s behavioral decision. Two weeks later the mice were tested again with the escape box in the original position (retention test). This allows for the examination of long-term memory. Finally, on the day after this test, the escape tunnel was moved to a new location (90 degrees from the original position) and the behavior of the mouse was recorded. This is called the reversal test and it allows for the examination of perseveration at the old hole as well as the working memory strategies the mice adopted to locate the new tunnel location. Each session was videotaped and scored by an experimenter blind to the genotype of the mouse. Measures recorded include the latency to enter the escape box and the number of errors made per session. Errors are defined as nose pokes and head deflections over any hole that did not have the tunnel beneath it. The probe data were analyzed using Noldus Ethovision software to determine time spent in each quadrant of the maze as well as to assess activity measures.

#### Novel Object Recognition

Mice were individually habituated to a 51cm x 51cm x 39cm open field for 5 min. Mice were then tested with two identical objects placed in the field (either two 250 ml amber bottles or two clear plastic cylinders 6×6×16cm half filled with glass marbles). An individual animal was allowed to explore for 5 min, now with the objects present. After two such trials (each separated by 1 minute in a holding cage), the mouse was tested in the object novelty recognition test in which a novel object replaced one of the familiar objects (for example, an amber bottle if the cylinders were initially used). All objects and the arena were thoroughly cleaned with 70% ethanol between trials to remove odors. Behavior was video recorded and then scored for number contacts (touching with nose or nose pointing at object and within 0.5 cm of object) and/or for time contacting the objects. Habituation to the objects across the familiarization trials (decreased contacts) was an initial measure of learning and then renewed interest (increased contacts) in the new object indicated successful object memory. Recognition indexes were calculated using the following formula: (# contacts during test)/(# contacts in last familiarization trial + # contacts during test). Values greater than 0.5 indicate increased interest, whereas values less than 0.5 indicate decreased interest in the object during the test relative to the final familiarization trial. quadrant of the maze as well as to assess activity measures.

#### Quantification and statistical analysis

All data were analyzed by t-test, 1-way, 2-way or 3-way between-subject ANOVA with post hoc comparisons depending on the number of variables and groups in each analysis. For ANOVA, if statistical significance (p < 0.05) was achieved, we performed post hoc analysis to account for multiple comparisons. The level of significance (a) was always set at 0.05. Survival curves were analyzed using Log-rank (Mantel-Cox) Test. Data were analyzed using Prism 7 (GraphPad Software, La Jolla, CA) and are represented as means and standard error of the means. All experiments and data analyses were conducted in a blinded fashion. All data were prepared for analysis with standard spread sheet software (Microsoft Excel).

## Acknowledgements

We thank members of the lab past and present, as well as former and current colleagues and collaborators for their helpful contributions.

