## Supplemental Figures for "Skeletal Muscle TFEB Signaling Promotes Central Nervous System Function and Reduces Neuroinflammation during Aging and Neurodegenerative Disease"

#### Supplemental Figure 1

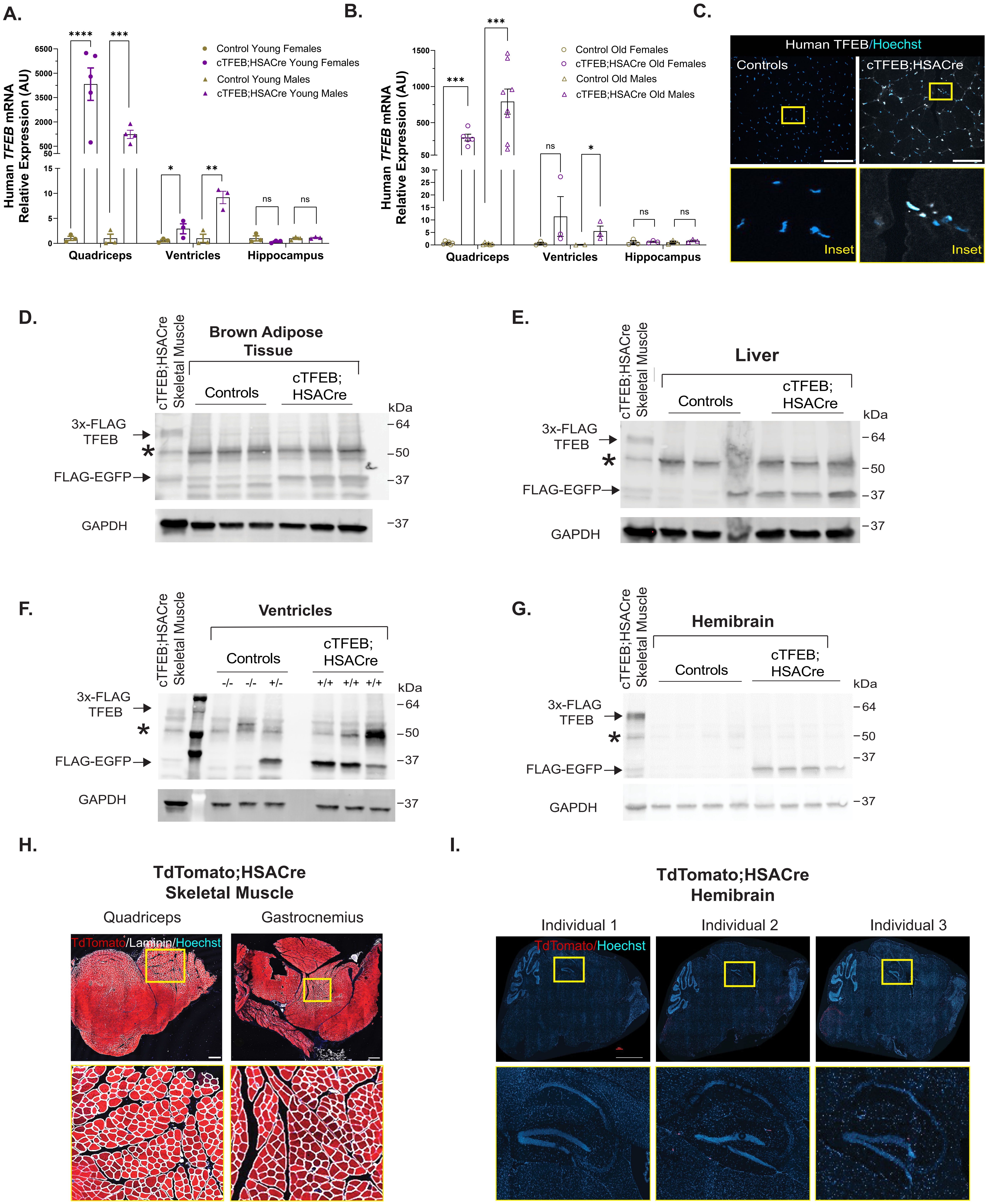

### Supplemental Figure 2

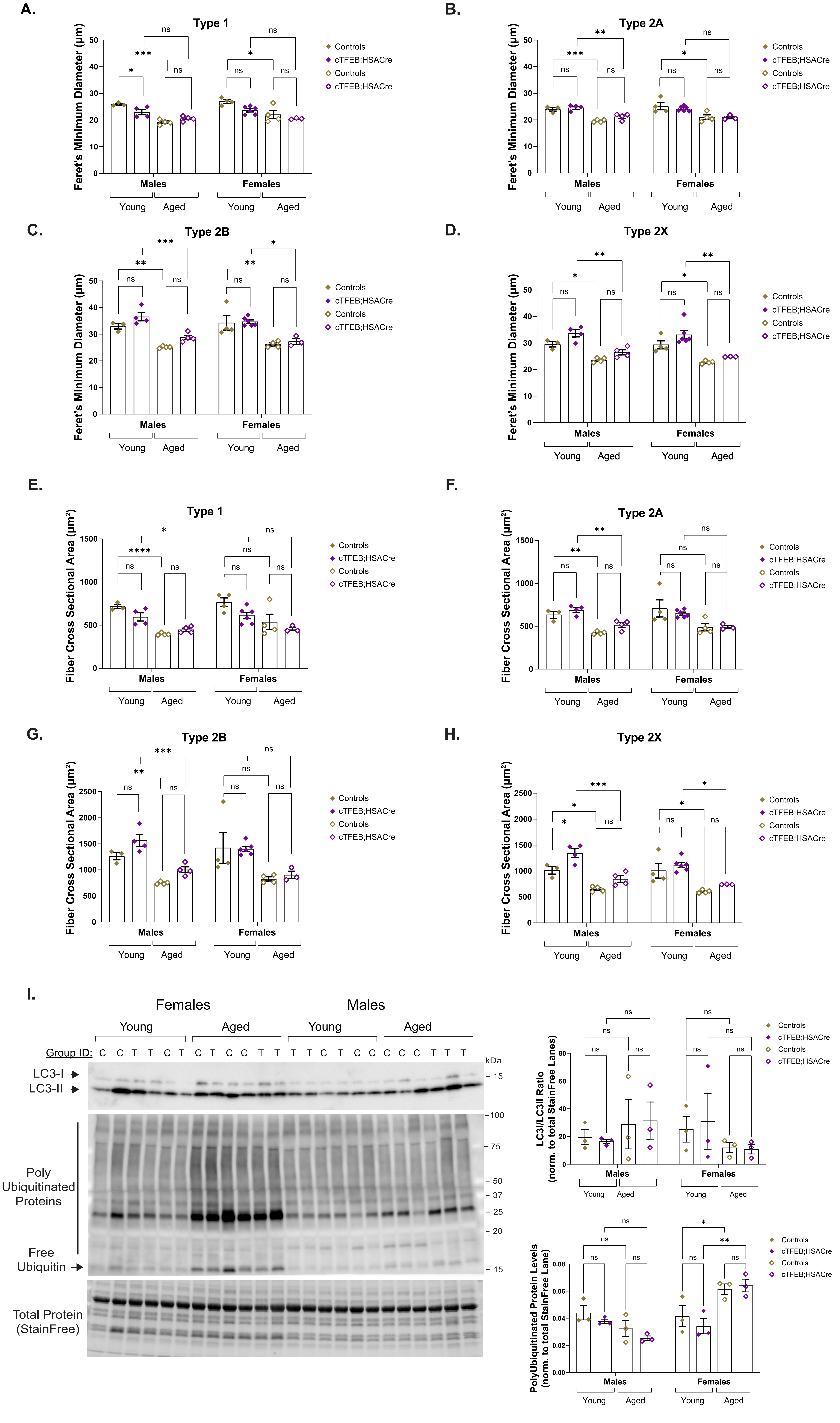

#### Supplemental Figure 3

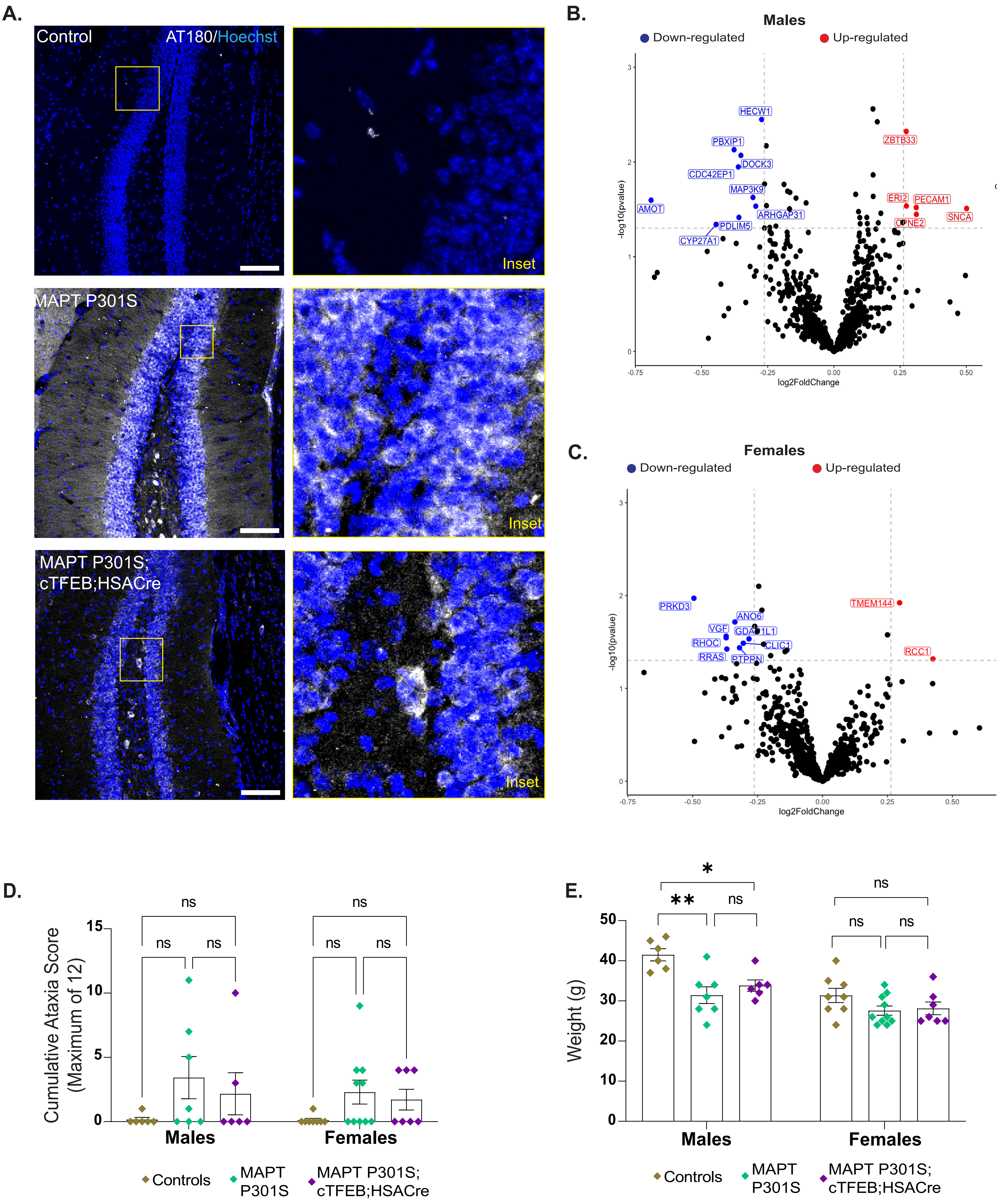

#### Supplemental Figure 4

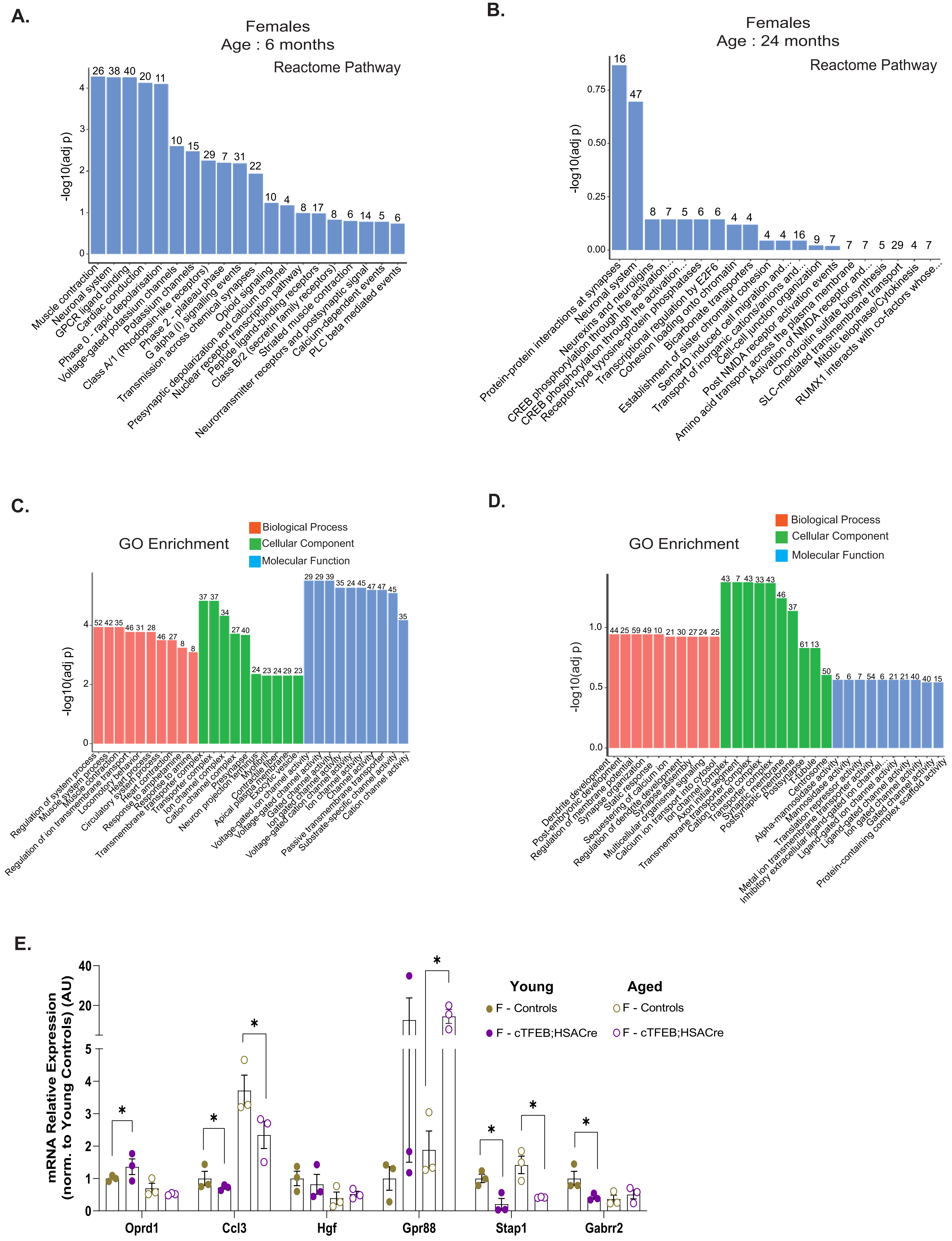

#### Supplemental Figure 5

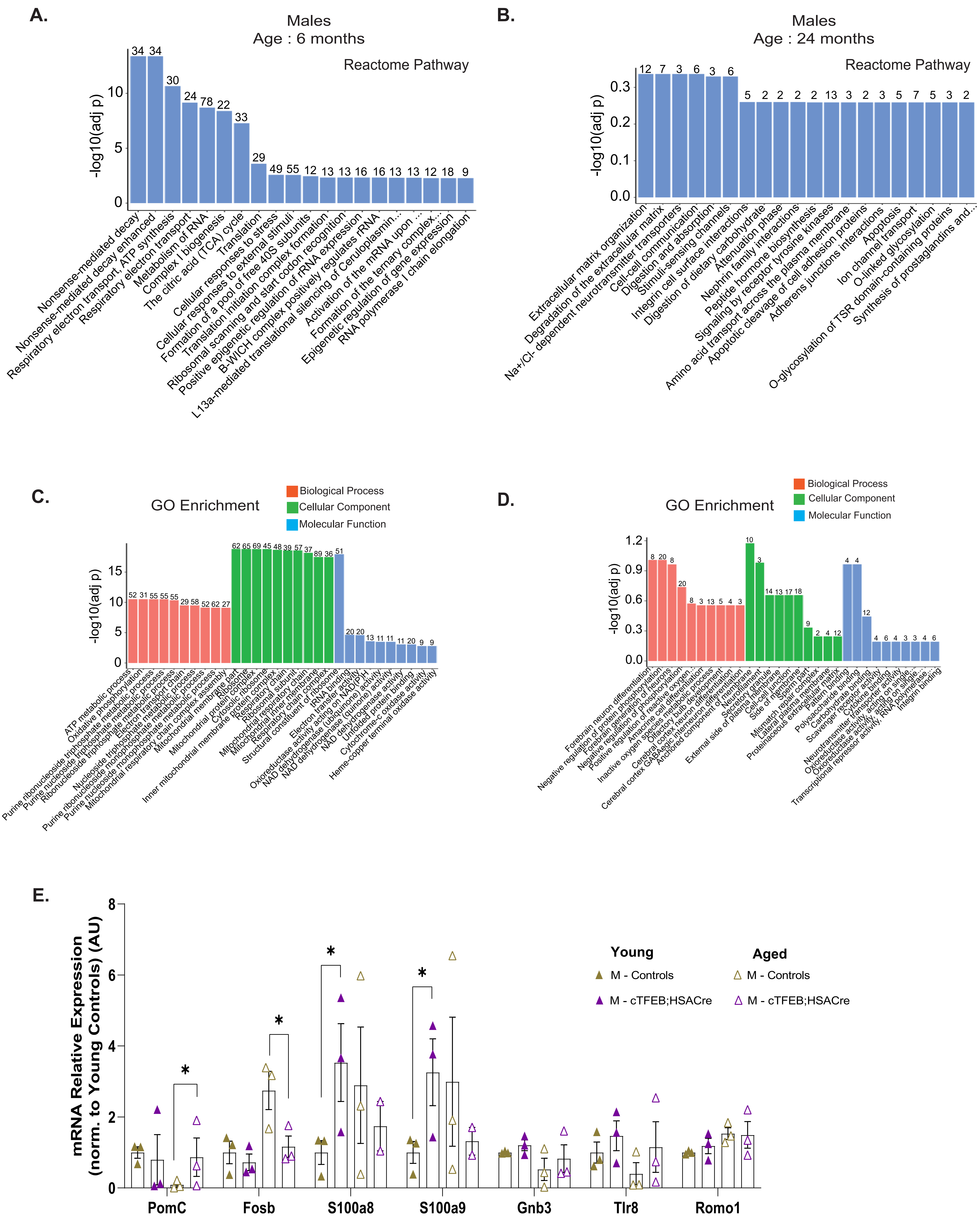

**Supplemental Figure 6**

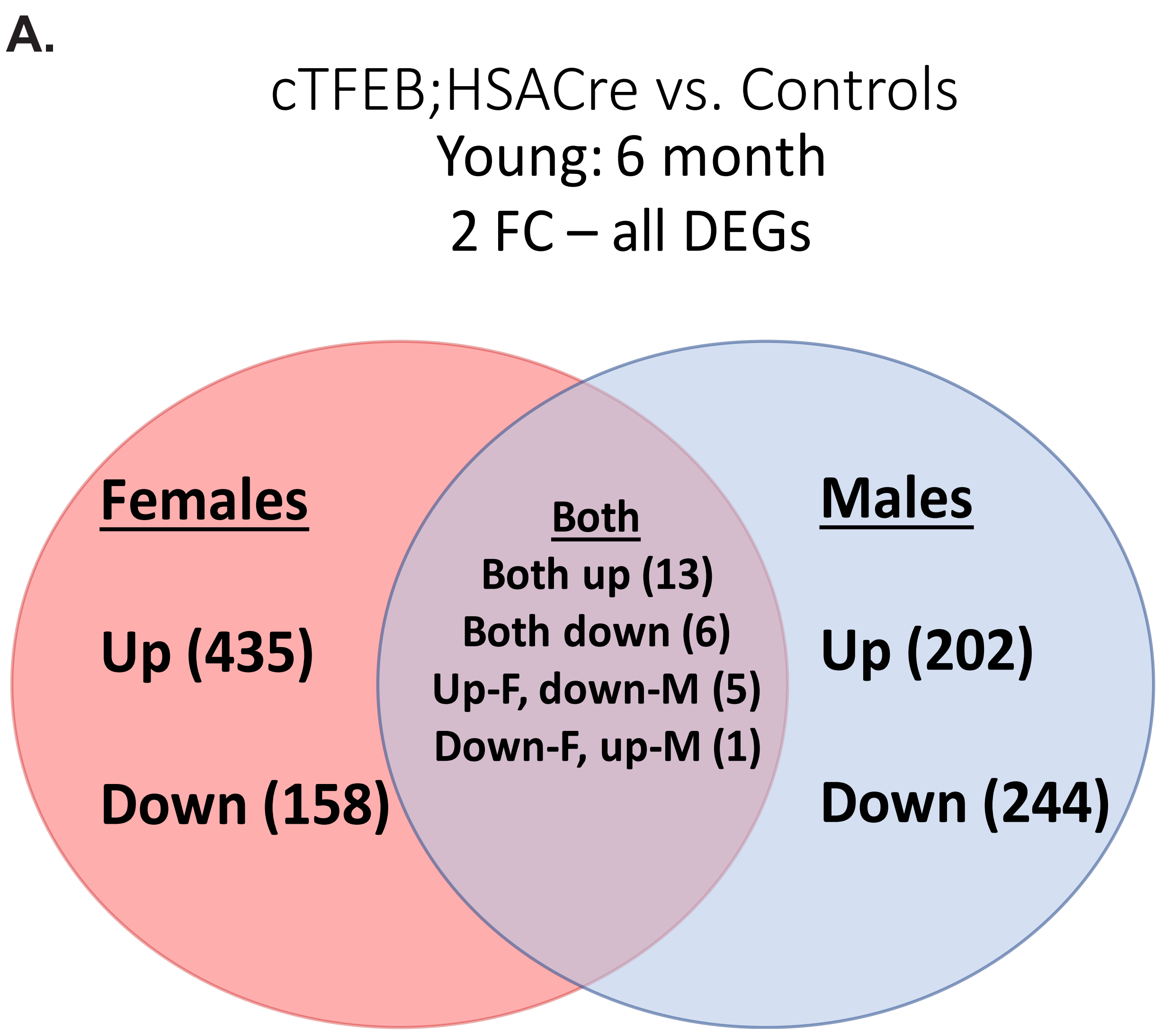


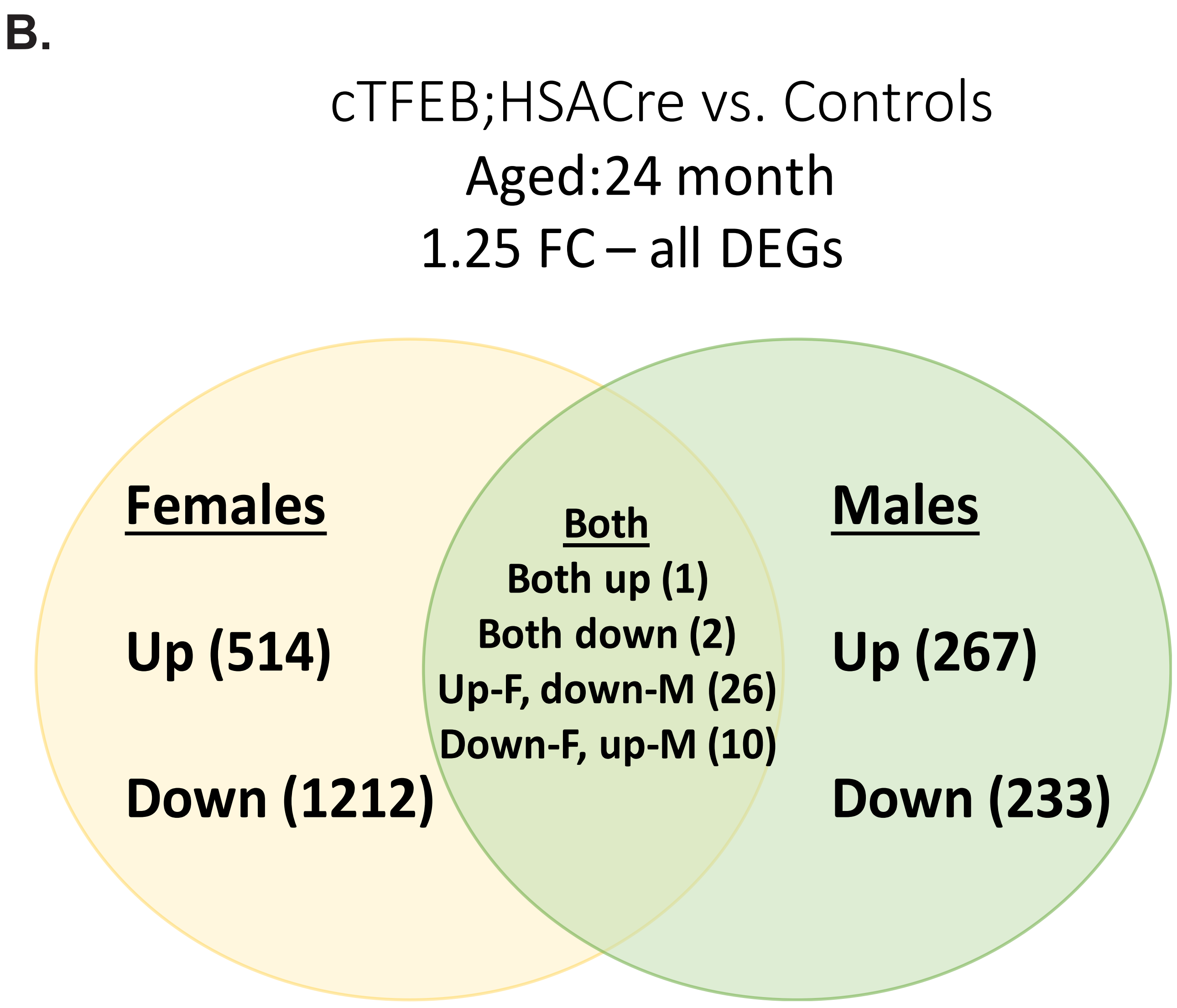
